# Membrane compartmentalization of Ect2/Cyk4/Mklp1 and NuMA/dynein/dynactin is essential for cleavage furrow formation during anaphase

**DOI:** 10.1101/2022.03.29.486209

**Authors:** Shrividya Sana, Ashwathi Rajeevan, Sachin Kotak

## Abstract

In animal cells, spindle elongation during anaphase is temporally coupled with cleavage furrow formation. Spindle elongation during anaphase is regulated by NuMA/dynein/dynactin complexes that occupy the polar region of the cell membrane and are excluded from the equatorial membrane. How NuMA/dynein/dynactin are excluded from the equatorial membrane and the biological significance of this exclusion remains unknown. Here, we show that the centralspindlin (Cyk4/Mklp1) and its interacting partner RhoGEF Ect2 are required for NuMA/dynein/dynactin exclusion from the equatorial cell membrane. The Ect2-based (Ect2/Cyk4/Mklp1) and NuMA-based (NuMA/dynein/dynactin) complexes occupy mutually exclusive membrane surfaces during anaphase. The equatorial membrane enrichment of Ect2-based complexes is essential for NuMA/dynein/dynactin exclusion and proper spindle elongation. Conversely, NuMA-based complexes at the polar region of the cell membrane ensure spatially confined localization of Ect2-based complexes and thus RhoA. Overall, our work establishes that membrane compartmentalization of NuMA-based and Ect2-based complexes at the two distinct cell surfaces restricts dynein/dynactin and RhoA for coordinating spindle elongation with cleavage furrow formation.

## Introduction

Animal cells elongate their mitotic spindle and segregate sister chromatids to the opposite poles before setting up their new boundary by forming a cleavage furrow during anaphase (reviewed in Green et al., 2012; Basant and Glotzer, 2018; Pollard and O’Shaughnessy, 2019). Spindle elongation and sister chromatids’ separation are tightly coordinated with cleavage furrow formation. The proper functioning of these processes is vital for preventing chromosomes instability and tumorigenesis (reviewed in Ganem et al., 2007; Lens and Medema, 2018).

Spindle elongation and chromosomes segregation is regulated by an evolutionarily conserved cortically anchored protein nuclear mitotic apparatus (NuMA) [reviewed in di Pietro et al., 2016; Bergstralh et al., 2017; Kotak, 2019; Lechler and Mapelli, 2021]. Cortically anchored NuMA serves as an adaptor for the microtubule-dependent minus-end-directed motor protein complex dynein and its associated dynactin complex (reviewed in Kiyomitsu, 2019; Kotak, 2019; Lechler and Mapelli, 2021). The pulling forces generated by the cortically anchored dynein/dynactin are assumed to promote spindle elongation and chromosomes segregation. The cortical level of NuMA during mitosis is temporally controlled by a biochemical cross-talk between cyclin-dependent kinase 1 (Cdk1) and PP2A-B55γ-based phosphatase complex (Kiyomitsu and Cheeseman, 2013; Kotak et al., 2013; Seldin et al., 2013; Zheng et al., 2014; Keshri et al., 2020). During anaphase, NuMA is localized to the cell cortex by directly associating with the membrane lipids PI(4)P and PI(4,5)P2 (referred to as PIP and PIP2) (Zheng et al., 2014; Kotak et al., 2014). Notably, despite the presence of PIP and PIP2 across the entire cell membrane, NuMA is excluded from the equatorial region of the cell membrane (Kotak et al., 2014). Further, it was shown the equatorial exclusion of NuMA is dependent on Rho GTPase-activating protein (RhoGAP) Cyk4 (also known as MgcRacGAP) (Kotak et al., 2014). However, the mechanism and the biological relevance of Cyk4-dependent NuMA exclusion from the equatorial membrane is not known.

In metazoans, the cleavage furrow formation at the equatorial membrane is initiated by local activation of the small GTPase RhoA. RhoA directly controls actin polymerization and indirectly regulates myosin II activation at the equatorial membrane and thus helps in cleavage furrow formation. Coordinated assemblies of multiple protein complexes at the spindle midzone (also known as the central spindle) regulate the spatiotemporal activation of RhoA at the equatorial membrane (reviewed in Green et al., 2012; Basant and Glotzer, 2018; Pollard and O’Shaughnessy, 2019). The spindle midzone is a stable array of overlapping microtubules of opposite polarity that assembles during metaphase-to-anaphase transition halfway between the segregating chromosomes. One of the critical complexes that assemble at the spindle midzone and regulate RhoA activation at the equatorial membrane is centralspindlin (Somers and Saint, 2003; Yuce et al., 2005). Centralspindlin is a heterotetrametric complex consisting of a dimer of kinesin-6 family member mitotic kinesin-like protein (Mklp1), and a dimer of Cyk4 (Pavicic-Kaltenbrunner et al., 2007). The centralspindlin helps in recruiting the conserved RhoA guanine nucleotide exchange factor (RhoGEF)-epithelial cell transforming sequence 2 (Ect2) to the spindle midzone (Yuce et al., 2005; Chalamalasetty et al., 2006; Kamijo et al., 2006; Su et al., 2011; Kotynkova et al., 2016; Gomez-Cavazos et al., 2020; Schneid et al., 2021). Ect2 is essential for RhoA activation and thus for cleavage furrow ingression and cytokinesis in animal cells (Miki et al., 1993; Prokopenko et al., 1999; Kimura et al., 2000; Su et al., 2011; reviewed in Basant and Glotzer, 2017; Pollard and O’Shaughnessy, 2019). Ect2 consists of BRCT (BRCA1-C-terminal) domains at the N-terminus, and RhoGEF, pleckstrin homology (PH), and poly-basic cluster (PBC) domains at the C-terminus (Chalamalasetty et al., 2006; Su et al., 2011; Kotynkova et al., 2016; Schneid et al., 2021; Figure 3A). Through BRCT domains, Ect2 interacts with Cyk4, and the PH and PBC domains are critical for its membrane localization (Burkard et al., 2007; Somers and Saint, 2003; Yuce et al., 2005; Wolfe et al., 2009; Su et al., 2011; Kotynkova et al., 2016; Gomez-Cavazos et al., 2020; Schneid et al., 2021). The presence of these multiple domains in Ect2 ensures Ect2 localization and RhoA-activation at the equatorial region of the membrane for cleavage furrow formation. However, how Ect2 levels at the equatorial membrane are confined and maintained to a narrow region for proper RhoA activation is poorly understood and is a crucial aspect of understanding the molecular mechanism of cleavage furrow formation.

In this study, we show that polarized distribution of NuMA/dynein/dynactin (also referred to as NuMA-based complexes) at the polar region of the cell membrane and Ect2/Cyk4/Mklp1 (also referred to as Ect2-based complexes) at the equatorial region of the cell membrane are critical in ensuring proper spindle elongation, and timely cleavage furrow formation. In cells depleted for Ect2, or Cyk4, NuMA-based complexes are localized to the equatorial cell membrane, affecting chromosomes’ separation kinetics. Vice versa, in cells depleted for NuMA, the Ect2-based complexes fail to confine to the equatorial membrane, thus impacting RhoA accumulation. Our data further reveal that NuMA localization at the polar membrane acts redundantly with the spindle midzone localized Ect2-based complexes to ensure proper cleavage furrow formation. In summary, this work provides insights into the mechanism that limits RhoA and dynein/dynactin at two membrane surfaces by establishing mutually exclusive membrane localization of two evolutionarily conserved protein complex assemblies.

## Results

### Centralspindlin and Rho GEF Ect2 exclude NuMA from the equatorial membrane

During anaphase, NuMA is restricted to the polar region of the cell membrane, while the equatorial membrane is mutually exclusively occupied by RhoA (Figure 1A-1C). We showed earlier that siRNA-mediated depletion of Cyk4, which is critical for RhoA accumulation, results in ectopic accumulation of NuMA and its associated dynein/dynactin motor protein complexes at the equatorial membrane (Kotak et al., 2014). To investigate the mechanism of NuMA/dynein/dynactin exclusion from the equatorial membrane in anaphase, we first analyzed the localization of NuMA and dynactin subunit p150^Glued^ in cells depleted for proteins that act downstream of Cyk4 and are crucial for RhoA recruitment (Figure 1D; reviewed in Eggert et al., 2006; Green et al., 2012; Basant and Glotzer, 2018). HeLa cells transfected with siRNA against Cyk4, Mklp1, Ect2, or Anillin lead to efficient depletion of these proteins, and significant accumulation of bi-nucleated or multi-nucleated cells, indicating robust cytokinesis failure (Figure 1E-1I; Figure S1A-S1D). As expected, depletion of Cyk4 and its associated kinesin Mklp1 resulted in NuMA and p150^Glued^ localization at the equatorial membrane (Figure 1J-1N and 1Q). Further, cells transfected with Ect2 siRNA showed robust enrichment of NuMA and p150^Glued^ at the equatorial membrane (Figure 1O and 1Q).

**Figure 1.**
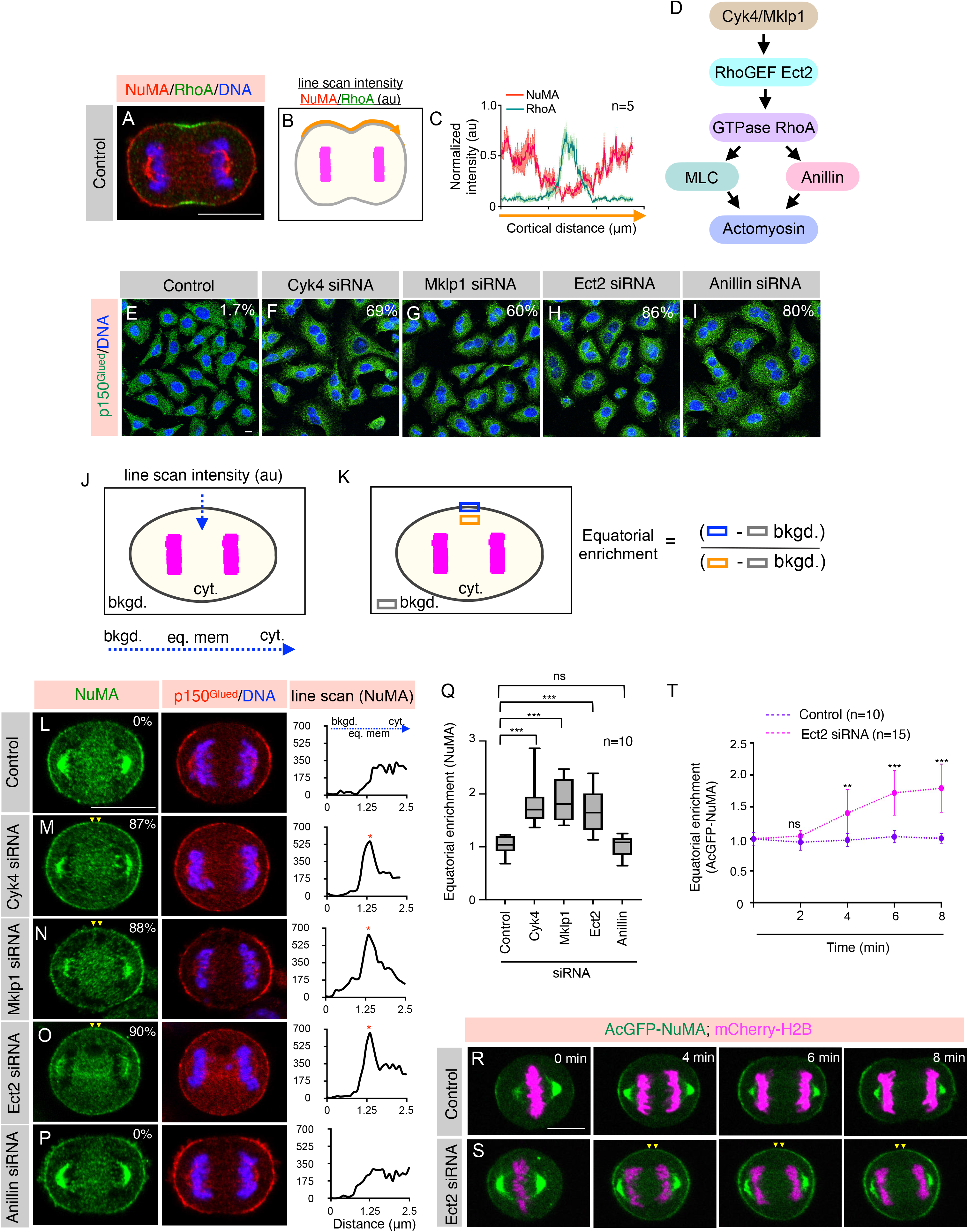
Centralspindlin (Cyk4/Mklp1), and Ect2 are required for NuMA/p150^Glued^ exclusion from the equatorial membrane. (A) Immunofluorescence (IF) analysis of HeLa cells in anaphase. Cells were fixed and costained using anti-RhoA (green), and anti-NuMA (red) antibodies, as indicated. In this, and other IF-analysis panels DNA is shown in blue unless specified. More than 30 cells were analyzed and the representative cells are shown here. Scale bar in this and for the following panels represent 10 µm. (B, C) Schematic representation of line scan analysis (B), and the outcome (C) of such analysis for RhoA and NuMA normalized intensity [in arbitrary unit (au)] for the specified membrane region (in orange). (n=5 cells were used for this quantification as depicted; shaded region indicates SEM). (D) Schematic representation of an evolutionarily conserved pathway required for RhoA activation in metazoans. (E-I) IF analysis of HeLa cells that are either transfected with control siRNA (E) or siRNA-against Cyk4 (F), Mklp1 (G), Ect2 (H), or Anillin (I). Cells were fixed after 36 hr of siRNA transfection and stained with anti-p150^Glued^ antibodies (green). Multinucleation percentage (%) is shown for each siRNA condition (n>500 cells each from three independent experiments). Knockdown efficiency was also determined by immunoblot analysis (see Figure S1). (J, K) Schematic representation of line scan analysis (J), and quantification method (K) used for analyzing equatorial membrane enrichment of NuMA fluorescence intensity. bkgd., eq. mem., and cyt. represent background, equatorial membrane and cytoplasmic intensity. (L-P) HeLa cells transfected with control siRNA (L) or siRNA against-Cyk4 (M), Mklp1 (N), Ect2 (O), or Anillin (P). Cells were fixed after 36 hr of siRNA transfection and thereafter costained with anti-NuMA (green), and anti-p150^Glued^ (red) antibodies. Percentage on the figure panels represent the fraction of anaphase cells that show equatorial NuMA enrichment for each condition as analyzed by visual quantification (n > 100 cells from three independent experiments). Yellow arrowheads depict NuMA localization at the equatorial membrane. Line scan analysis on the right represents NuMA fluorescence intensity as indicated in Figure panel J. Red asterisk represents intensity at the equatorial membrane. (Q) Quantification of equatorial membrane NuMA intensity in cells transfected with control siRNA or siRNA against-Cyk4, Mklp1, Ect2, or Anillin as indicated in Figure panel K (n=10 cells for all conditions, as listed). In this and other Figures ns-p0.05; *-p<0.05; **-p<0.01; ***-p<0.001 as determined by two-tailed Student’s t-test; error bars: SD). (R, S) Confocal live-imaging analysis of HeLa Kyoto cells stably coexpressing AcGFP-NuMA (green) and mCherry-H2B (magenta) that are transfected with control siRNA (R) or siRNA against Ect2 (S). Recording was started 26 hr post transfection for control and Ect2 siRNA. ’0 min’ time-point represent metaphase to anaphase transition. Yellow arrowheads depict NuMA localization at the equatorial membrane. (T) Quantification of equatorial membrane AcGFP-NuMA intensity as described in Figure panel K (n=10 cells for control siRNA, and n=15 for Ect2 siRNA transfected cells; error bars: SD).

We next assessed the importance of other proteins that act downstream of RhoA for NuMA and p150^Glued^ exclusion from the equatorial membrane. Depletion of proteins such as Anillin-a structural component of the cytokinetic contractile ring, and mDia (also known as Diaphanous-related formin-1)-a protein required for actin polymerization at the cytokinetic ring, did not cause NuMA and p150^Glued^ mislocalization to the equatorial membrane (Figure 1P and 1Q; Figure S1E-S1H). Since RhoA is essential for myosin II activation (Figure 1D; reviewed in Green et al., 2012; Basant and Glotzer, 2018; Pollard and O’Shaughnessy, 2019), we next asked if myosin II activity is crucial in excluding NuMA from the equatorial membrane. For this purpose, we utilized a non-photo toxic myosin II inhibitor para-nitroblebbistatin (PNBB; Kepiro et al., 2014) and treated mitotically synchronized HeLa cell population with PNBB (Figure S1I). Despite a strong impact of PNBB on myosin II activity that resulted in a significant increase in bi-nucleated cells (Figure S1J and S1K), NuMA remained excluded from the equatorial membrane in cells treated with PNBB during anaphase (Figure S1L-S1O).

To track the spatiotemporal localization of NuMA, we performed live-cell imaging in monoclonal stable HeLa Kyoto cells coexpressing AcGFP (Aequora coerulescens GFP) and a mono FLAG epitope-tagged NuMA and mCherry-H2B (Rajeevan et al., 2020). Analogous to endogenous protein, AcGFP-NuMA localizes to the equatorial membrane in cells transfected with Ect2 siRNA, compared to control cells (Figure 1R-1T). Similar results were seen upon Cyk4 or Mklp1 depletion (data not shown). Since NuMA is a cortical adaptor for dynein, we further analyzed dynein localization in cells stably expressing GFP-tagged dynein heavy chain (DHC1) (Poser et al., 2008). We found that similar to NuMA localization, GFP-DHC1 is no longer restricted to the polar region of the cell cortex in cells transfected with Ect2 siRNA (Figure S1P and S1Q). Altogether, these results suggest that the centralspindlin complex (Cyk4/Mklp1) and Ect2 are necessary for excluding NuMA/dynein/dynactin from the equatorial membrane.

However, the contractile ring proteins which acts downstream of RhoA and that we tested in our study, do not appear to be essential for NuMA exclusion from the equatorial membrane.

### Failure in cell elongation upon Ect2 depletion is not responsible for NuMA exclusion

Cells depleted either for the centralspindlin complex or Ect2 fail to elongate significantly during anaphase. Therefore, an alternative possibility could be that the compromised cell elongation in cells that are depleted for Ect2 (or Cyk4/Mklp1) is the cause for NuMA/dynein/dynactin localization at the equatorial membrane. To determine whether it is (1) cells’ inability to elongate upon centralspindlin/Ect2 depletion or (2) a direct role of these proteins in NuMA exclusion, we monitored the equatorial localization of NuMA in control and Ect2 siRNA transfected cells at early stages of anaphase before cell elongation begins. We noticed no significant difference in the cell length in cells transfected with Ect2 siRNA versus control siRNA at 1-4 min post anaphase entry (Figure 2A-2D). However, cells transfected with Ect2 siRNA significantly localized AcGFP-NuMA at the equatorial membrane as early as 3 min post anaphase entry (Figure 2A-2D).

**Figure 2.**
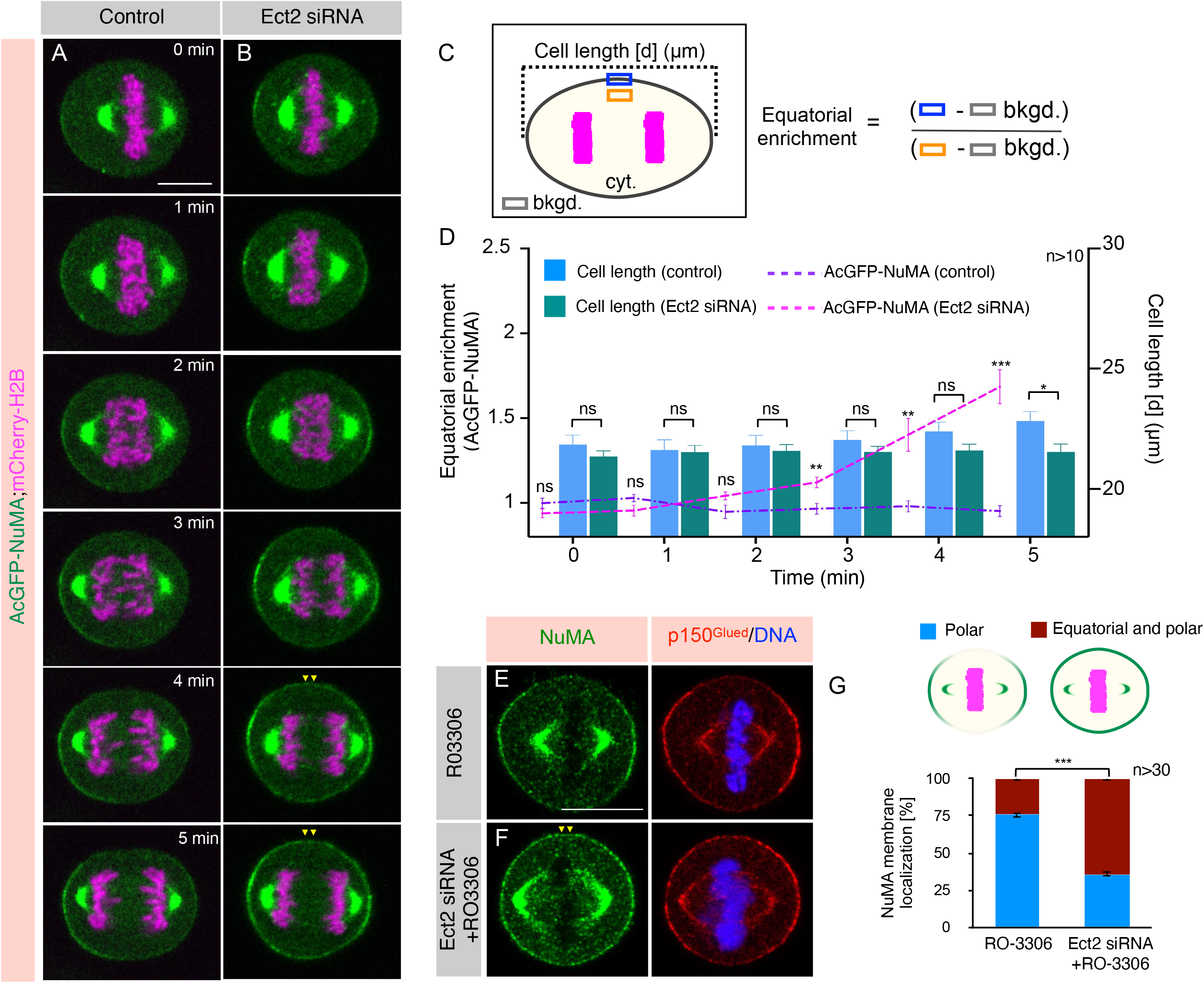
NuMA/p150Glued localization at the equatorial membrane is not because of cell elongation failure. (A, B) Confocal live-imaging analysis of HeLa Kyoto cells stably coexpressing AcGFP-NuMA (green) and mCherry-H2B (magenta) that are transfected with control siRNA (A) or siRNA against Ect2 (B). Recording was started 26 hr post transfection for control and Ect2 siRNA transfected cells. ‘0 min’ time-point represent metaphase to anaphase transition, and the images were acquired every minute. Yellow arrowheads depict NuMA localization at the equatorial membrane. Scale bar in this and for the following panels represent 10 µm. (C, D) Schematic representation of quantification method used for analyzing equatorial membrane enrichment of AcGFP-NuMA fluorescence intensity and for computing cell length [d (µm)] (C). Quantification of equatorial membrane enrichment of NuMA, along with the cell length (D) as described pictorially in Figure panel C. Bar graph represents the cell-length for cells that are either transfected with control siRNA (blue), or siRNA against Ect2 (green). Line plot graph represents the equatorial NuMA membrane intensity in cells either transfected with control siRNA (violet) or Ect2 siRNA (pink). Time (in min) is shown on the x-axis and NuMA equatorial intensity together with the cell length is plotted on the y-axis. (n>10 cells as listed, error bars: SEM). (E, F) IF analysis of HeLa cells that were either control siRNA transfected and treated with RO-3306 (E), or transfected with siRNA-against Ect2 and treated with RO-3306 (F) for 5 min. Cells were fixed and costained using anti-NuMA (green), and anti-p150^Glued^ (red) antibodies. Yellow arrowheads depict NuMA localization at the equatorial membrane. (G) Visual quantification of the cortical NuMA distribution in RO-3306, or in Ect2 siRNA plus RO-3306 treated cells. Stacked blue bar, and brown bar represent the percentage of the cells that show cortical NuMA distribution at the polar cortical region, and equatorial and polar cortical region respectively (n>30 cells, from three independent experiments; error bars: SD).

**Figure 3.**
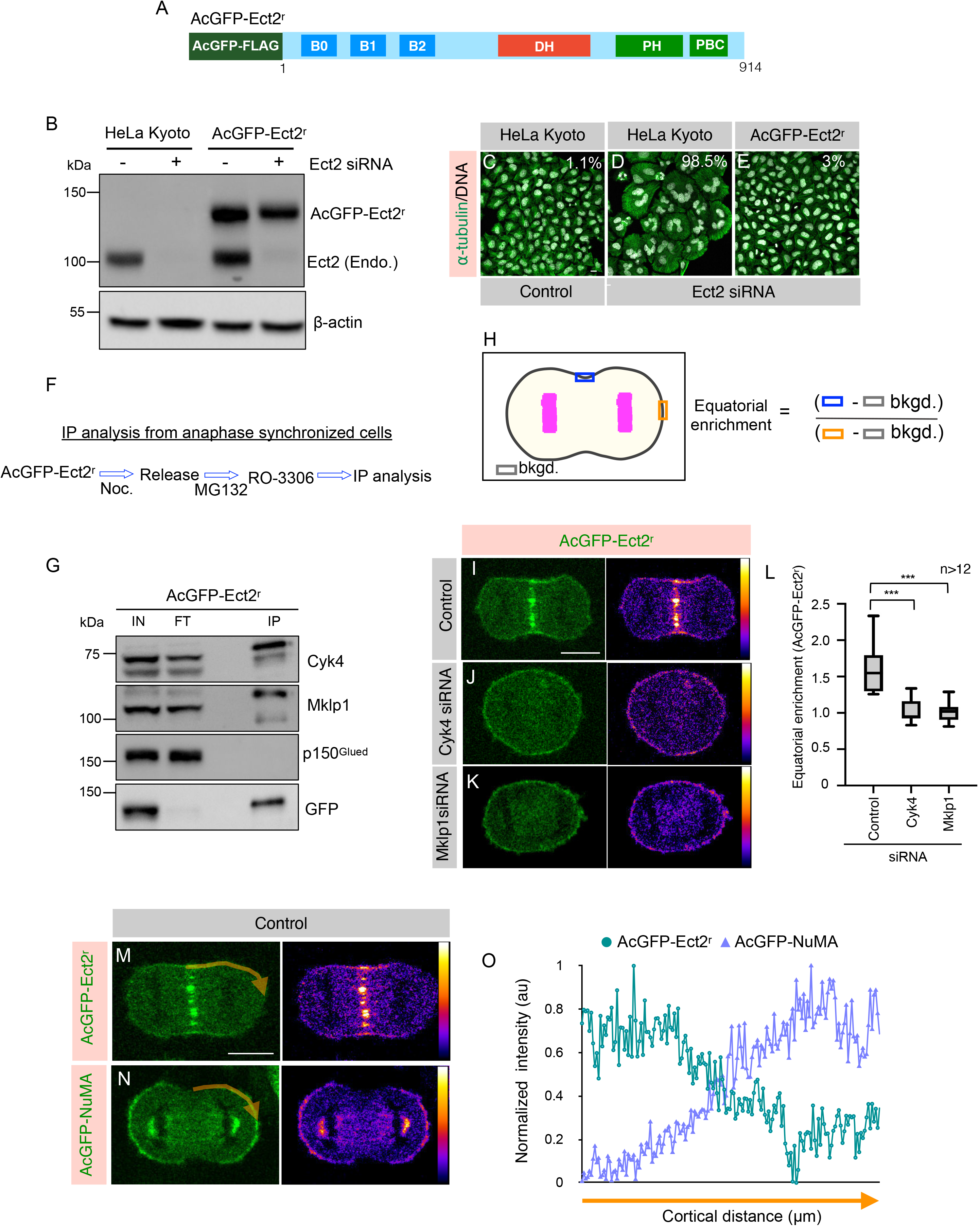
Ect2/Cyk4/Mklp1-based complexes are localized to the equatorial membrane in anaphase. (A) Schematic representation of siRNA-resistant Ect2 construct with AcGFP (*Aequora coerulescens* GFP)-tag, and mono FLAG-tag at the N-terminus (referred to as AcGFP-Ect2^r^). (B) Immunoblot analysis of mitotically synchronized protein extracts made from HeLa Kyoto cells, or Kyoto cells that were stably expressing AcGFP-Ect2^r^. These extracts were prepared 60 hr post transfection with control siRNA (-), or siRNA against Ect2 (+). Resulting blot was probed with antibodies directed against Ect2 and β-actin. As mentioned transgenic AcGFP-Ect2^r^ and endogenous Ect2 were detected on this immunoblot. In this and other panels, the molecular mass is indicated in kilodaltons (kDa) and is shown on the left. (C-E) IF analysis of the Kyoto cells that are transfected with control siRNA (C), siRNA against Ect2 (D) or Kyoto cells stably expressing AcGFP-Ect2^r^ after transfection with siRNA against Ect2 (E) for 60 hr post transfection. Cells were fixed and stained using anti-⍺-tubulin (green) antibody. DNA is shown in grey. Multinucleation percentage (%) is shown for each siRNA condition. (n>500 cells each from three independent experiment). Scale bar in the panel and following panels represent 10 µm. (F) Experimental protocol for chemically induced anaphase onset of mitotically synchronized HeLa Kyoto cells stably expressing AcGFP-Ect2^r^ by acute Cdk1 inactivation using RO-3306. (G) Co-immunoprecipitation (IP) by GFP-Trap from chemically induced anaphase synchronized mitotic HeLa Kyoto cells lysates stably expressing AcGFP-Ect2^r^. Resulting blots were probed for Cyk4, Mklp1, p150^Glued^, and GFP as indicated. IN (1% of total), IP: 20% of the total. Please note that for GFP detection in the IP fraction, only 3% of IP fraction was loaded. Note that AcGFP-Ect2^r^ interacts with Cyk4, and Mklp1 (centralspindlin complex), but not with dynein interacting dynactin subunit p150^Glued^. (H) Schematic representation of quantification method used for analyzing equatorial membrane enrichment of AcGFP-Ect2^r^ fluorescence intensity. (I-K) Images from confocal live-imaging analysis of HeLa Kyoto cells stably expressing AcGFP-Ect2^r^ (green) that are transfected with control siRNA (I) or siRNA against Cyk4 (J), or Mklp1 (K). Signal intensity of AcGFP-Ect2^r^ is also specified by pseudocolor gradient on the right side of the image. (L) Equatorial membrane quantification for AcGFP-Ect2^r^ for control siRNA, Cyk4 siRNA, or Mklp1 siRNA was performed as described in the Figure panel H. (n>12 cells, error bars: SD). (M-O) The membrane localization of AcGFP (green) in HeLa Kyoto cells that are either expressing AcGFP-Ect2^r^ (M) or AcGFP-NuMA (N). Signal intensity of AcGFP-Ect2^r^ or AcGFP-NuMA is also specified by pseudocolor gradient on the right side of the image. The linescan analysis of the membrane AcGFP intensity for the region (in orange) is depicted in O. For such measurements one-quadrant of an anaphase cell from the equatorial cell membrane to the polar region of the cell membrane was analyzed for calculating the normalized intensity (O).

To further corroborate that the equatorial membrane localization of NuMA is possibly not because of the altered cell size in Ect2 depleted cells, we analyzed the localization of NuMA and p150^Glued^ in roundish metaphase cells that are acutely treated with Cdk1 inhibitor RO-3306 (Vassilev et al., 2006), to program them in an anaphase like state. Brief incubation (5 min) of metaphase cells with RO-3306 facilitates centralspindlin accumulation at the spindle midzone as observed by analyzing Cyk4 localization (Figure S2A and S2B; Keshri et al., 2020), and in majority (75%) of these cells NuMA is excluded from the equatorial membrane (Figure 2E and 2G). However, a significant fraction of metaphase cells that are transfected with Ect2 siRNA and are treated with RO-3306 robustly localize NuMA at the equatorial membrane (compare Figure 2F with 2E; Figure 2G). These results strongly suggest that NuMA localization in the equatorial membrane in Ect2 depleted cells is not simply because of failure in proper cell elongation.

### Chromosome passenger complex (CPC) is dispensable for NuMA exclusion from the equatorial membrane

Centrosome-localized Aurora A kinase phosphorylates NuMA and regulates its cortical distribution in metaphase (Gallini et al., 2016; Kotak et al., 2016). Therefore, we wondered if Aurora B, an essential part of chromosome passenger complex (CPC) which localizes to the spindle midzone and regulates cytokinesis, is involved in excluding NuMA from the equatorial membrane (Gruneberg et al., 2004; Hummer and Mayer, 2009; Kitagawa et al., 2013; Kitagawa et al., 2014; Basant et al., 2015). To this end, we combined Aurora B inactivation with cell synchronization using a specific Aurora B inhibitor, ZM447439 (Figure S3A; Ditchfield et al., 2003). Despite significant cytokinesis failure in cells that are treated with ZM447439, NuMA/p150^Glued^ remained restricted to the polar region of the membrane identical to the control cells during anaphase (Figure S3B-S3E). To strengthen these findings, we further depleted kinesin-6 Mklp2, which is necessary to localize Aurora B at the spindle midzone in anaphase (Figure S3F-S3I; Gruneberg et al., 2004; Hummer and Mayer, 2009; Kitagawa et al., 2013). siRNA-mediated depletion of Mklp2 did not impact cortical NuMA/p150^Glued^ distribution (compare Figure S3K with S3J). These results indicate that CPC component Aurora B which localizes to the central spindle similar to the centralspindlin complex is not involved in NuMA/dynactin exclusion from the equatorial membrane.

### NuMA exclusion from the equatorial membrane is crucial for efficient chromosome separation

NuMA/dynein/dynactin localization at the polar cortical region during anaphase is critical for proper spindle elongation, possibly by generating cortical pulling forces (Collins et al., 2012; Kotak et al., 2013; Zheng et al., 2014; Keshri et al., 2020; reviewed in Kotak, 2019; Kiyomitsu and Boerner, 2021). Thus, we set out to determine whether the presence of NuMA/dynein/dynactin at the equatorial membrane in cells depleted for Ect2 or centralspindlin component Cyk4 perturbs proper spindle elongation due to exerting a counteracting force from the equatorial cortical region. To this end, we performed the live-imaging analysis in cells coexpressing AcGFP-NuMA and mCherryH2B and are transfected with either control or Ect2 siRNA. Since spindle elongation is coupled with chromosome separation in human cells (Roostalu et al., 2010), we measured the distance between separating sister chromatids in cells undergoing metaphase-to-anaphase transition. We noticed a modest but significant (p <0.001) impact on the chromosome separation kinetics in cells depleted for Ect2 (Figure S4A and S4B). Similar results were obtained in cells transfected with Cyk4 siRNA (data not shown). These data indicate that the ectopic accumulation of NuMA/dynein/dynactin at the equatorial membrane may result in an imbalance of pulling forces that hamper proper chromosomes segregation kinetics and possibly spindle elongation during anaphase.

### Ect2/Cyk4/Mklp1 tripartite complex excludes NuMA from the equatorial membrane

To get a mechanistic insight on how Ect2, Cyk4, and Mklp1 can exclude NuMA, we generated monoclonal stable HeLa Kyoto cells expressing siRNA-resistant allele of Ect2 that is N-terminally tagged with AcGFP and a FLAG-tag (referred to as AcGFP-Ect2^r^), similar to what has been reported previously (Figure 3A; Su et al., 2011). This transgenic line expresses ectopic AcGFP-Ect2^r^ protein in amounts comparable to an endogenous copy of the gene (Figure 3B). The robust cytokinesis failure seen upon the depletion of endogenous Ect2 is fully rescued in this cell line (Figure 3B-3E), suggesting that the AcGFP-Ect2^r^ protein is functional. Immunoprecipitates (IP) from anaphase-synchronized cell extract made from AcGFP-Ect2^r^ expressing cells revealed that AcGFP-Ect2^r^ interacts with endogenous Cyk4 and Mklp1, as reported previously (Figure 3F and 3G; Somers and Saint, 2003; Yuce et al., 2005). This data suggests Cyk4, Mklp1, and Ect2 make a tripartite complex in anaphase. AcGFP-Ect2^r^, which is in complex with Cyk4 and Mklp1 during anaphase, is confined to the equatorial membrane, and this restricted localization of AcGFP-Ect2^r^ is significantly perturbed in cells depleted for Cyk4 or Mklp1 (Figure 3H-3L). Crucially, AcGFP-Ect2^r^ localizes to the equatorial membrane surfaces devoid of AcGFP-NuMA (Figure 3M-3O). These data indicate that Ect2 enriches at the equatorial membrane in a complex with centralspindlin (Cyk4/Mklp1) and raise a possibility that the confined localization of Ect2/Cyk4/Mklp1-based tripartite complex at the equatorial membrane may restrict NuMA/dynein/dynactin to the polar membrane (see discussion).

How might Ect2/Cyk4/Mklp1-based tripartite complex exclude NuMA/dynein/dynactin from the equatorial cell membrane? Previous work established that carboxy-terminal of Ect2 (Ect2CT) possess a conserved pleckstrin homology (PH) and polybasic cluster (PBC) that is capable of directly interacting with [PI(4)P], and [PI(4,5)P2] (Chalamalasetty et al., 2006; Su et al., 2011). Notably, NuMA C-terminus has been shown to interact with identical phosphoinositides (Kotak et al., 2014; Zheng et al., 2014). Therefore, we sought to investigate if the localization of Ect2/Cyk4/Mklp1-based tripartite complex at the equatorial membrane via membrane-binding potential of Ect2 is responsible for NuMA/dynein/dynactin exclusion. To this end, we generated a monoclonal HeLa Kyoto cell line expressing siRNA-resistant allele of Ect2 lacking its C-terminal PH and PBC domain and tagged with AcGFP and a FLAG-tag (Figure 4A; and referred to as AcGFP-Ect2^r^). The transgenic protein made by AcGFP-Ect2^r^_Δmem_ expressing cell line is analogous to AcGFP-Ect2^r^ expressing line (Figure 4B). Depletion of endogenous protein in cells expressing wild-type allele of Ect2 (AcGFP-Ect2^r^), but not AcGFP-Ect2^r^_Δmem_ rescue cytokinesis failure as reported earlier (Figure 4C-4F; Su et al., 2011). Next, we examined the localization of NuMA in these lines upon endogenous Ect2 protein depletion. Transiently expressed mCherry-NuMA is excluded from the equatorial membrane in cells stably expressing AcGFP-Ect2^r^, and AcGFP-Ect2^r^ is localized to the spindle midzone and at the proximal equatorial membrane in these cells (Figure 4G, 4I and 4J). As reported previously, AcGFP-Ect2^r^_Δmem_ localizes at the spindle midzone, but fails to accumulate at the equatorial cell membrane (Figure 4H; Su et al., 2011). Importantly, mCherry-NuMA localizes to the equatorial membrane surface in cells stably expressing AcGFP-Ect2^r^_Δmem_ (Figure 4H, 4I and 4J). These results suggest that the membrane-binding ability of Ect2 ensures the enrichment of Ect2/Cyk4/Mklp1 complexes at the equatorial membrane, which helps in the exclusion of NuMA/dynein/dynactin complex from the equatorial membrane.

**Figure 4.**
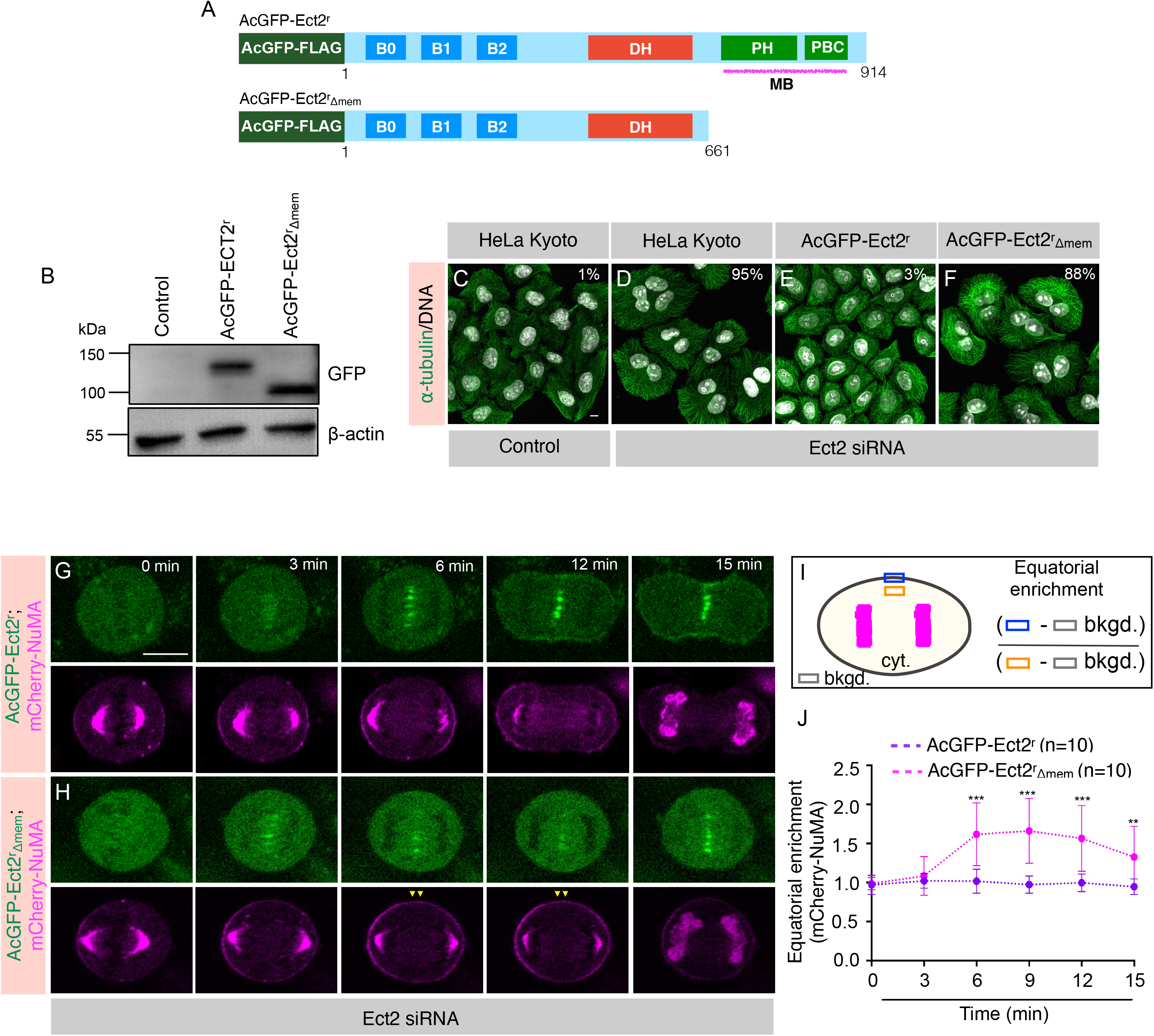
Membrane binding region of Ect2 is essential for NuMA exclusion from the equatorial membrane. (A) Schematic representation of AcGFP (*Aequora coerulescens* GFP), and mono FLAG -tag siRNA-resistant Ect2 full-length (referred to as AcGFP-Ect2^r^), and Ect2 without its membrane binding (PH and PBC) domain (referred to as AcGFP-Ect2^r^_Δmem_). (B) Immunoblot analysis of mitotically synchronized protein extracts made from HeLa Kyoto cells, Kyoto cells that were stably expressing either AcGFP-Ect2^r^ or AcGFP-Ect2^r^_Δmem_. Resulting blot was probed with anti-GFP or anti-β-actin antibodies. (C-F) IF analysis of the HeLa Kyoto cells transfected with either control siRNA (C), siRNA against Ect2 (D), HeLa Kyoto cells stably expressing AcGFP-Ect2^r^ and transfected with Ect2 siRNA (E), or HeLa Kyoto cells stably expressing AcGFP-Ect2^r^_Δmem_ and transfected with Ect2 siRNA (F) for 48 hr. Cells were fixed and stained with anti-⍺-tubulin (green) antibody. DNA is shown in grey. % on each panel marks the cytokinesis failure as calculated by analyzing the occurrence of binucleated and multinucleated cells (n>500 cells each from three independent experiments). Scale bar in the panel and following panels represent 10 µm. (G, H) Confocal live-imaging analysis of HeLa Kyoto cells stably expressing AcGFP-Ect2^r^ (G), or AcGFP-Ect2^r^_Δmem_ (H), which are transiently transfected with mCherry-NuMA and are depleted for endogenous Ect2 by siRNA. Recording was started 32 hr post transfection for control and Ect2 siRNA targeting endogenous Ect2. ‘0 min’ time-point represent metaphase to anaphase transition. Yellow arrowheads depict NuMA localization at the equatorial membrane. (I) Schematic representation of quantification method used for analyzing equatorial membrane enrichment of mCherry-NuMA fluorescence intensity. (J) Equatorial membrane quantification for mCherry-NuMA in cells expressing either AcGFP-Ect2^r^, or AcGFP-Ect2^r^_Δmem_ and are depleted for endogenous Ect2 as described in the above figure panel I. (n=10 cells; error bars: SD)

### NuMA at the polar membrane ensures proper RhoA levels at the equatorial membrane

Our results indicate that the enrichment of Ect2/Cyk4/Mklp1 complexes at the equatorial membrane prevent NuMA/dynein/dynactin accumulation at those membrane surfaces. In that case, conversely, loss of NuMA from the polar region of the membrane should allow Ect2-based complexes to spread to the distant membrane, and concomitantly, this should lead to a reduction in the levels of these complexes at the equatorial membrane. To assess the localization of Ect2-based tripartite complexes in cells depleted of NuMA, we analyzed the localization of RhoA since Ect2-based complexes are critical for regulating and confining RhoA at the equatorial membrane (Yuce et al., 2005). Cells transfected with NuMA siRNA show a modest but significant reduction in RhoA levels at the equatorial membrane during anaphase in comparison to control cells (compare Figure 5B with 5A, quantification in Figure 5E-5J). However, despite this significant reduction in RhoA levels at the equatorial membrane, cells transfected with NuMA siRNA establish and ingresses cytokinetic furrow at a time comparable to control cells (Figure 6B; Movie S2). We reasoned that no impact of NuMA depletion on cytokinetic furrow could be due to robust enrichment of Ect2/Cyk4/Mklp1 complexes at the spindle midzone, which would rapidly exchange with the proximal equatorial membrane and will maintain a critical RhoA level, even when NuMA is not present at the polar membrane. What would happen to cleavage furrow if the localization of Ect2/Cyk4/Mklp1 complexes at the spindle midzone is compromised along with NuMA depletion? To test this possibility, we analyzed the localization of RhoA in cells codepleted for NuMA and protein regulator of cytokinesis 1 (Prc1). Prc1 cross-links antiparallel microtubules and is vital for the assembly of the spindle midzone (Figure S5A-S5C; Mollinari et al., 2002; Mollinari et al., 2005; Zhu et al., 2006; Kellogg et al., 2016). Prc1 depletion drastically diminishes Cyk4, Mklp1, and Ect2 localization at the spindle midzone (Figure 5K-5R). However, Prc1 depletion does not significantly impact RhoA levels at the equatorial membrane, as reported previously (compare Figure 5C with 5A; quantification in Figure 5E-5J; Verbrugghe and White, 2004; Mollinari et al., 2005). Importantly, we observed either significantly diminished or no RhoA enrichment at the equatorial membrane in cells codepleted for NuMA and Prc1 (Figure 5DcatA, 5DcatB, and related quantification in Figure 5E-5J). RhoA enrichment zone that usually occupies ∼12% of the cell perimeter during mid-anaphase in control cells (Figure S5D and S5E) had possibly spread to the distant membrane in cells codepleted for NuMA and Prc1 (Figure 5G and 5H), which likely resulted in reduced intensity of RhoA at the equatorial membrane. Also, we confirmed that this diminished RhoA zone in cells codepleted for NuMA and Prc1 is not because of any direct impact of NuMA and Prc1 depletion on the total protein levels of centralspindlin component Cyk4, Ect2, or RhoA during mitosis (Figure 5S).

**Figure 5.**
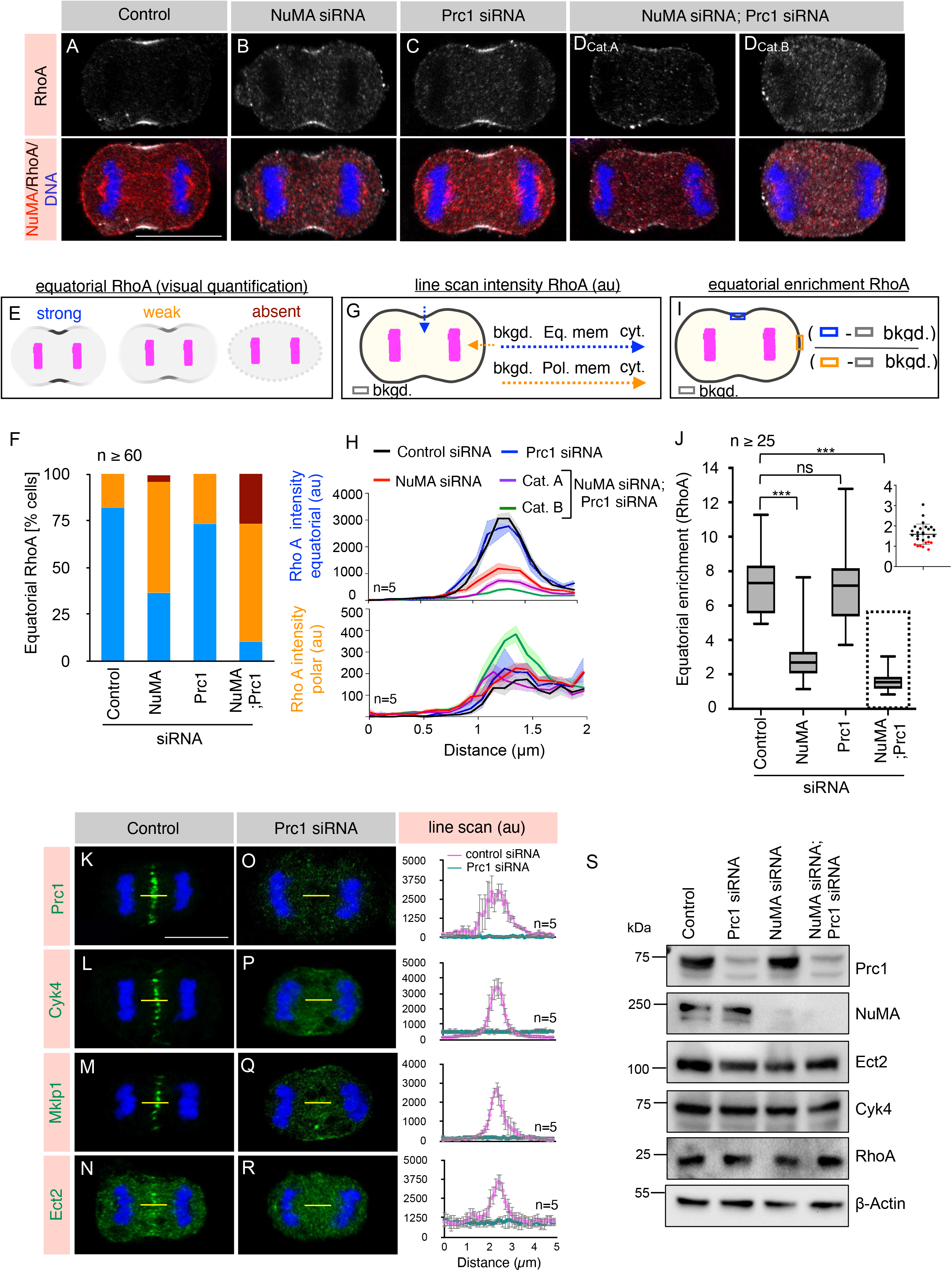
NuMA and Prc1 codepletion impact RhoA accumulation at the equatorial membrane. (A-D) Immunofluorescence (IF) analysis of HeLa Kyoto cells that are either transfected with control siRNA (A) or siRNA-against NuMA (B), Prc1 (C), or Prc1 and NuMA [D_cat. A_ and D_cat. B_]. Cells were fixed 60 hr post siRNA transfection and stained with anti-RhoA (grey) and anti-NuMA (red) antibodies. A minimum of 60 anaphase cells from three independent experiments were analyzed visually and representative images are shown in the Figure panel. Scale bar in the panel and following panels represent 10 µm. (E, F) Schematic representation of the visual quantification of equatorial RhoA membrane intensity (E), and the % of cells based on such visual quantification in various siRNA transfected conditions (F). For such analysis, cells were grouped in three categories: strong (depicted in blue), as in control siRNA transfected cells, weak (depicted in orange), as in NuMA siRNA, and absent (in brown), as in NuMA and Prc1 siRNA. (n=109 cells for control siRNA; n=81 cells for NuMA siRNA; n=96 cells for Prc1 siRNA, and n=60 cells for NuMA and Prc1 siRNA transfected cells from three independent experiments). (G, H) Schematic representation of line scan analysis (G), and the outcome (H) of such analysis for RhoA intensity at the equatorial membrane (depicted in blue), and polar membrane (depicted in orange) in different siRNA transfected conditions. Please note a decrease in RhoA equatorial intensity in NuMA siRNA, or NuMA and Prc1 siRNA transfected cells. Also, note a marginal increase in RhoA intensity at the polar membrane in cells transfected with NuMA and Prc1 siRNA. (n=5 representative cells from all siRNA conditions were used for this quantification; shaded region indicates SEM) (I, J) Schematic representation of the quantification method (I), and the outcome of such analysis for equatorial membrane enrichment of RhoA fluorescence intensity (J). Please note significant decrease in equatorial membrane RhoA intensity in cells depleted for NuMA, or NuMA and Prc1. The distribution for the RhoA intensity for all the cells depleted for NuMA and Prc1 is shown in the inset. Note approximately 30% of cells show the ratio of a selected region at the equatorial membrane to that of a similar area at the polar membrane is ∼1 (shown as red dots in the inset). (n=25 cells for control siRNA, NuMA siRNA or Prc1 siRNA, and n=27 cells for NuMA and Prc1 siRNA transfected cells; error bars: SD). (K-R) Line scan analysis of the central spindle localization of Prc1 (K, O), Cyk4 (L, P), Mklp1 (M, Q), or Ect2 (N, R) in HeLa Kyoto cells that are transfected with either control siRNA or siRNA against Prc1. Yellow-line represent the area that was used for computing the line-scan intensity (in arbitrary unit, au) on the right of each panel. (n=5 cells in each condition, as depicted; error bars: SD). (S) Immunoblot analysis of mitotically synchronized protein extracts made from HeLa Kyoto cells that were either transfected with control siRNA, or siRNA against Prc1, NuMA, or Prc1 and NuMA for 60 hr. Resulting blot was probed with anti-NuMA, Anti-Prc1, anti-Ect2, anti-Cyk4, and anti-RhoA antibodies. Anti-β-actin antibodies were used to analyze the equal loading of the samples.

**Figure 6.**
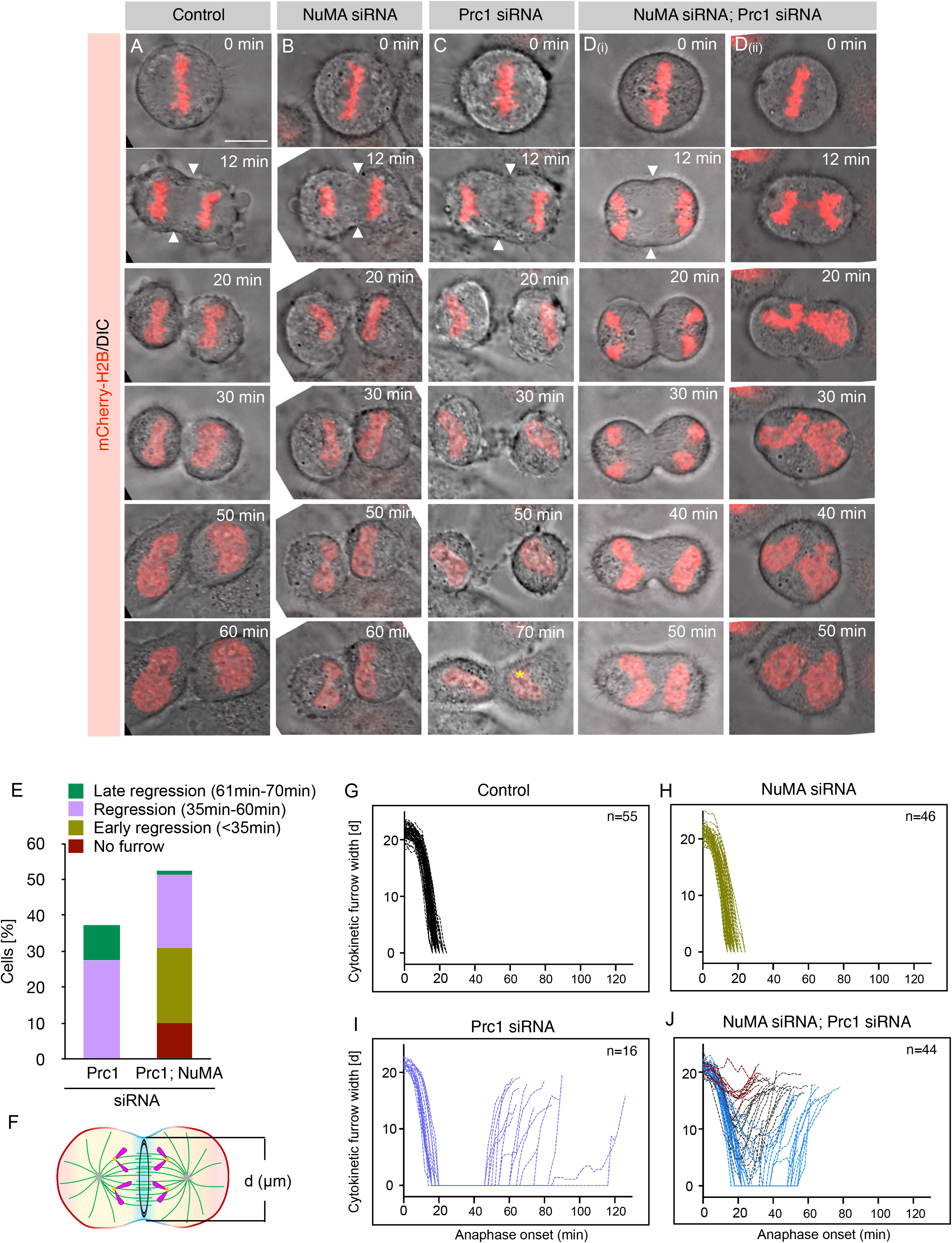
Co-depletion of NuMA and Prc1 affect cytokinetic furrow initiation. (A-D) Confocal live-imaging analysis in combination with Differential Interphase Contrast (DIC) microscopy images of HeLa Kyoto cells stably expressing mCherry-H2B, and were either transfected with control siRNA (A), siRNA against NuMA (B), siRNA against Prc1 (C), or siRNA against Prc1 and NuMA (D(i) and D(ii)). D(i) represents cells where the cytokinetic furrow was initiated but do not fully ingress, and regresses back. While D(ii) represents cells where no cytokinetic furrow initiation was observed. Recording was started 50 hr post transfection. ‘0 min’ time-point represent metaphase to anaphase transition. White arrowhead represents appearance of cytokinetic furrow. Yellow asterisk depicts cytokinesis failure (binucleated cell) in Prc1 siRNA transfected cell after complete furrow ingression, followed by regression. Scale bar in these panels represent 10 µm. (E) Quantification of various cytokinetic phenotypes as indicated in HeLa Kyoto cells stably expressing mCherry-H2B that were transfected with Prc1 siRNA or Prc1 and NuMA siRNA (n>50 cells). Recording was started 50 hr post transfection ‘0 min’ time-point represent metaphase to anaphase transition, and for every cell time-lapse movies were recorded till 70 min from time 0 min. Since control siRNA transfected cells, and NuMA siRNA transfected cells do not reveal any furrow regression, and cytokinesis failure within 70 min of time-lapse movies acquisition, their data are not included in the graph. (Total cells analyzed for Prc1 siRNA and for NuMA and Prc1 siRNA transfected cells were 51 and 87 respectively) (F-J) Schematic representation of anaphase cell depicting the method of measurements of the cytokinetic furrow width [d (µm)] from time-lapse confocal live imaging obtained for panel A-D (F). Furrow width measurements in cells stably expressing mCherry-H2B that were transfected with control siRNA (G), siRNA against NuMA (H), siRNA against Prc1 (I), and siRNA against Prc1 and NuMA (J). Please note that in contrast to Prc1 siRNA transfected cells, 10% of cells transfected with NuMA and Prc1 siRNA do not initiate cytokinetic furrow ingression, while 20% of these cells do not fully ingress cytokinetic furrow, and proceed with furrow regression in time less than 35 min from time ‘0 min’ that indicates metaphase to anaphase transition. (a minimum of 16 cells were used for such analysis for each siRNA condition, as depicted on the Figure panels).

In sea urchin embryos, astral microtubules are linked in maintaining localized RhoA zone at the equatorial membrane (Bement et al., 2005; Foe and von Dassow, 2008, von Dassow et al., 2009). Thus, we wondered if the impact of Prc1 and NuMA codepletion on RhoA localization could be via their influence on the astral microtubules. However, we found no significant difference in the intensity profile of astral microtubules in cells transfected with siRNA against Prc1 and NuMA in comparison to the control cells (Figure S5F and S5G). Altogether these data strongly indicate that NuMA’s polarized membrane distribution acts in concert with the spindle midzone to ensure confined RhoA accumulation at the equatorial membrane, possibly by excluding Ect2/Cyk4/Mklp1-based tripartite complexes from the polar membrane surface.

### NuMA-based complexes at the polar membrane act redundantly with the spindle midzone to control cleavage furrow ingression

To scrutinize the relevance of the diminished or weak RhoA zone for cytokinesis in cells depleted for NuMA and Prc1, we conducted live-imaging experiments in HeLa Kyoto cells stably expressing mCherry-H2B. Control cells initiate cytokinetic furrow at ∼12 min and completely ingress furrow within ∼20 min (Figure 6A, 6F and 6G; Movie S1). As mentioned previously, despite a significant reduction in RhoA levels at the equatorial membrane in cells transfected with NuMA siRNA, these cells do not show any sign of cytokinesis defects (Figure 6B, 6F and 6H; Movie S2). As expected from the RhoA equatorial levels, Prc1 depletion in cells does not impact the timing of furrow initiation, but invariably these cells fail cytokinesis because of the instability of the abscission zone (Figure 6C, 6E, 6F and 6I; Movie S3). Notably, ∼20% of cells co-depleted for NuMA and Prc1 start furrow ingression at a time comparable to the control cells (∼12 min); however, the cleavage furrow does not fully ingress and undergoes regression, leading to cytokinesis failure (Figure 6D(i), 6E, 6F and 6J; Movie S4). 10% of NuMA and Prc1 co-depleted cells do not initiate furrow and completely fail cytokinesis (Figure 6D(ii), 6E, 6F and 6J; Movie S5). Taken together, these results suggest that the compartmentalized distribution of NuMA at the polar region of the cell membrane act redundantly with the Ect2/Cyk4/Mklp1 accumulation at the spindle midzone to maintain a confined zone of RhoA for proper assembly and ingression of cleavage furrow.

## Discussion

Polarized distribution of protein complexes at the cell membrane is critical for polarity establishment, spindle positioning, and cytokinesis. How protein complexes regulating these processes are localized and maintained at different membrane regions remains incompletely understood. During anaphase, antiparallel microtubules emanated from the two opposite centrosomes are dynamically reorganized between the segregated sister chromatids and form a spindle midzone (reviewed in Green et al., 2012; Avino et al., 2015). The assembly of multiple-protein complexes at the spindle midzone and the proximal membrane establishes a narrow RhoA zone at the equatorial membrane for cleavage furrow formation (reviewed in Bement et al., 2006; Green et al., 2012; Basant and Glotzer, 2018). Yet, how membrane RhoA levels are confined and maintained to the narrow equatorial zone of the cell membrane to promote cytokinesis has been unclear. It is hypothesized that a cross-talk between a stimulatory cue at the equatorial membrane and an inhibitory signal at the polar region of the cell cortex maintains RhoA to a restricted membrane zone (reviewed in Green et al., 2012; Basant and Glotzer, 2018). However, the molecular nature of inhibitory signal/s and the mechanism that prevents RhoA accumulation at the polar region of the cell membrane remains largely unknown. Our data suggest that the polarized distribution of NuMA-based complexes (NuMA/dynein/dynactin) at the polar region of the cell membrane restricts the localization of RhoA, possibly via antagonizing with Ect2-based complexes (Ect2/Cyk4/Mklp1) at the equatorial membrane. This is crucial for establishing a confined RhoA zone (Figure 7). We propose that the polarized distribution of NuMA/dynein/dynactin-based complexes could function as an inhibitory signal for RhoA accumulation by restricting Ect2/Cyk4/Mklp1 localization to the equatorial membrane. Conversely, the enrichment of Ect2-based complexes at the equatorial membrane limits NuMA/dynein/dynactin to the polar region of the cell membrane to ensure proper chromosomes separation (Figure S4).

**Figure 7.**
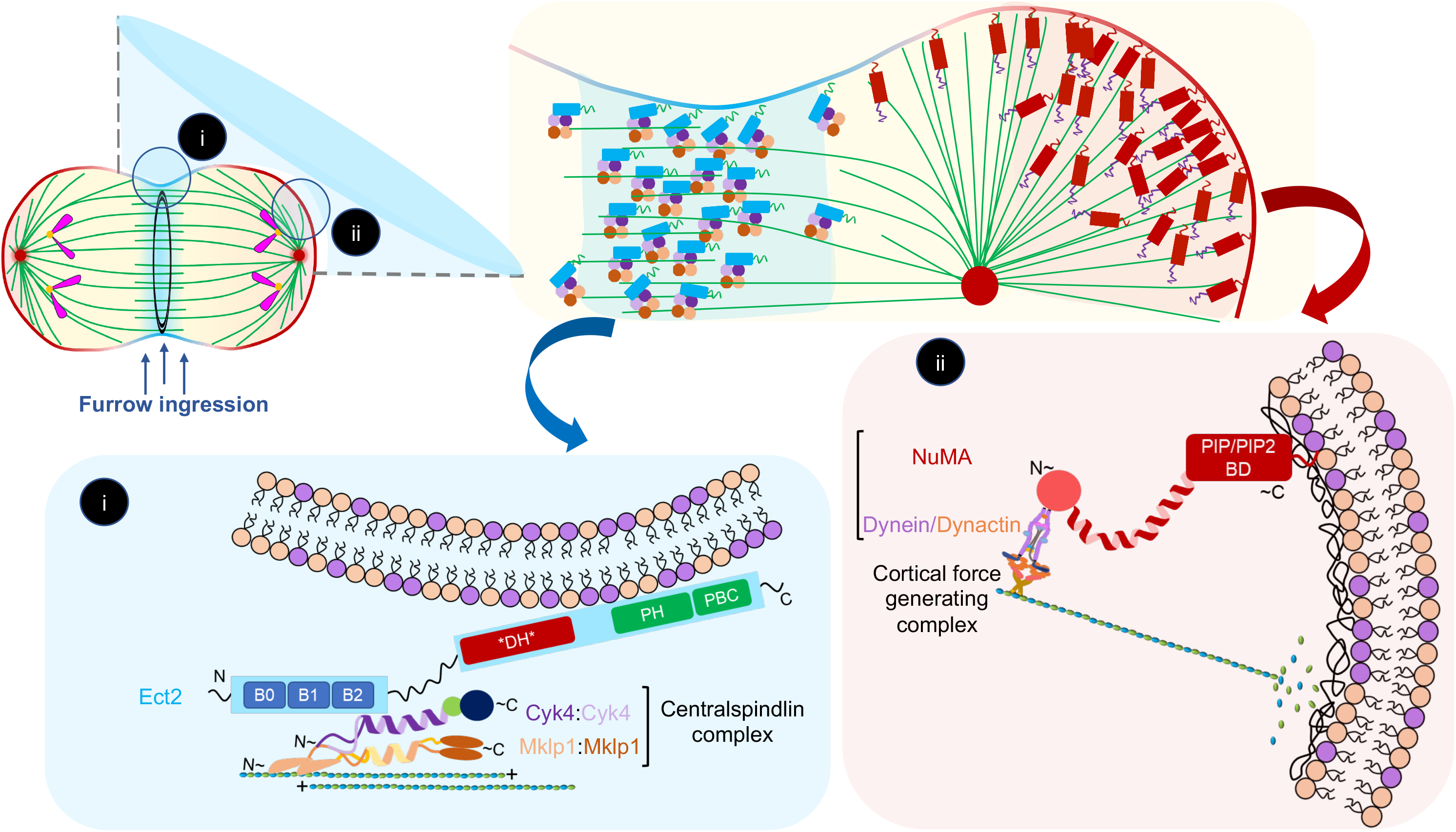
Ect2/Cyk4/Mklp1 and NuMA/dynein/dynactin-based complexes mutually antagonize, ensuring proper polarization and cytokinesis. Model for cortical localization of Ect2/Cyk4/Mklp1 and NuMA/dynein/dynactin at the equatorial cortical region and polar cortical region respectively. During anaphase assembly of spindle midzone ensure efficient localization of centralspindlin (2Cyk4:2Mklp1) together with its interacting component Ect2 enrichment at the central spindle, and at the equatorial cortical region. On the contrary NuMA/dynein/dynactin complexes enriches at the spindle poles, and at the polar cortical region. Inset i highlights the equatorial cortical region where Ect2 because of its ability to directly associate with the plasma membrane ensure constant flux of Ect2/Cyk4/Mklp1-based complexes to enrich at the equatorial surface, and regulate sufficient levels of RhoA for proper cytokinesis. In the absence of such robust flux [for instance in Prc1 (siRNA)], RhoA localization and cleavage furrow ingression is not affected because of the occupancy of NuMA/dynein/dynactin complexes at the polar region of the membrane (inset ii). This polarization between Ect2/Cyk4/Mklp1 and NuMA/dynein/dynactin ensure proper chromosomes separation, and the formation of cytokinetic furrow.

### Ect2-based complexes at the equatorial membrane restrict NuMA localization to the polar membrane

We uncovered that Ect2 and centralspindlin restrict NuMA/dynein/dynactin to the polar membrane. What is the mechanism of limiting NuMA/dynein/dynactin to the polar membrane? As established previously, we show that Ect2 is in a complex with Cyk4 and Mklp1 during anaphase (Somers and Saint, 2003; Yuce et al., 2005; Figure 3). This association ensures robust accumulation of Ect2-based complexes at the spindle midzone and the proximal equatorial membrane because of the membrane-binding potential of Ect2 (Figure 7i). We propose that enrichment of these Ect2-based complexes at the equatorial membrane creates spatial crowding that blocks NuMA/dynein/dynactin localization (Figure 7i). Indeed cells expressing Ect2Δmem, which can localize to the spindle midzone because of its interaction with centralspindlin but unable to localize at the membrane, fail to restrict NuMA at the polar region of the membrane.

Since membrane localization of Ect2-based complexes play a key role for RhoA activation and subsequently for contractile ring assembly, at this stage, we cannot rule out a possibility that the contractile ring protein/s other than Anillin or mDia (formin) could function either cooperatively with Ect2/Cyk4/Mklp1 or by yet unknown means to exclude NuMA/dynein/dynactin from the equatorial membrane. Moreover, small GTPase RhoA could also be directly involved in excluding NuMA-based complexes at the equatorial membrane. Since Ect2 and RhoA are engaged in a positive feedback loop (Chen et al., 2019), our results do not exclude an additional possibility that RhoA either alone or in combination with Ect2-based complexes is responsible for NuMA/dynein/dynactin exclusion. Nonetheless, our data support that myosin II activity is not critical for NuMA exclusion at the equatorial membrane during anaphase.

### NuMA localization at the polar membrane restricts the RhoA zone

NuMA/dynein/dynactin accumulates at the polar region of the membrane (Figure 7ii). We reasoned if the spatiotemporal localization of NuMA-based complexes at the polar membrane surface is due to spatial crowding created of Ect2-based complexes at the equatorial membrane, then NuMA depletion should allow Ect2-based complexes to spread to the distant membrane region. Based on this assumption, we should observe a spread in RhoA membrane signal and concomitantly decreased RhoA intensity at the equatorial membrane. Notably, cells depleted of NuMA alone had a significantly weak RhoA intensity at the equatorial membrane (Figure 5B). However, this reduction in RhoA levels did not impact cytokinetic furrow formation. This data aligns with the observations that furrow initiation requires significantly less RhoA activity (Loria et al., 2012; Tse et al., 2012). We rationalized that cells with NuMA depletion would still have sufficient RhoA levels at the equatorial membrane for cytokinetic furrow formation. This weak RhoA levels possibly exist because of the unperturbed pool of Ect2-based complexes at the spindle midzone, which would rapidly accumulate to the proximal equatorial membrane. If this positive regulatory arm is also removed, then we would expect to see further loss of RhoA levels at the equatorial membrane. Indeed, affecting the enrichment of Ect2-based complexes at the spindle midzone (by Prc1 siRNA) together with NuMA depletion further reduces the levels of RhoA intensity at the equatorial membrane. Notably, in a significant number of cells that were codepleted for NuMA and Prc1 (∼30%), the RhoA intensity at the equatorial membrane was highly reduced and in these cells the weak RhoA levels were detected at the polar membrane (Figure 5H). Since NuMA and Prc1 codepletion is not directly impacting RhoA, centralspindlin, or Ect2 protein levels in mitosis, we propose that reduction in RhoA intensity at the equatorial membrane in NuMA, and NuMA and Prc1 codepleted cells are simply because of more spread of RhoA regulators Ect2/Cyk4/Mklp1 to a larger surface area which is leading to an overall decrease in RhoA intensity at the equatorial membrane. We further noticed either a complete failure in furrow ingression, or ingression followed by regression in a significant number of cells codepleted for NuMA and Prc1 (Figure 6). These phenotypes were not seen in cells depleted for either NuMA or Prc1.

Our model of NuMA dependent physical exclusion of Ect2 based complexes from the polar cortex can also explain why depletion of Prc1 alone does not affect RhoA levels at the equatorial membrane and therefore has no impact on cytokinetic furrow formation. We reasoned that despite of the fact that depletion of Prc1 results in loss of Ect2-based complexes at the spindle midzone, their critical threshold at the equatorial membrane is constantly maintained owing to the presence of NuMA-based complexes at the polar membrane, which prevents them from diffusing out of the equatorial zone. This restricted mobility of Ect2-based complexes ensure timely cleavage furrow formation in Prc1 depleted cells (Figure 6; Verbrugghe and White, 2004; Mollinari et al., 2005).Therefore, we propose that NuMA-based complexes act redundantly with the spindle midzone localized Ect2-based complexes to establish a confined zone of RhoA for cleavage furrow formation. In summary, this work has characterized an essential but previously unrecognized mechanism that coordinates chromosome segregation with cleavage furrow formation by polarizing two evolutionarily conserved protein complexes at different regions of the cell membrane.

## Materials and methods

### Cell culture, transfections, and stable cell line generation

All the cell lines used in this study were cultured in high-glucose DMEM supplemented with 10% fetal bovine serum in a humidified 5% CO2 incubator at 37°C. For plasmid transfections, cells were seeded at 80% confluency in an imaging dish (Eppendorf, 0030740017) or on coverslips in six-well plates. Plasmid DNA (4 μg) suspended in 400 μl of serum-free DMEM was incubated for 5 min, followed by the addition of 6 μl of lipofectamine 2000 (Life Technologies, 11668019) or 6 μl turbofect (ThermoFisher Scientific, R0531), and was mixed and incubated for 15 min. This mixture was then added onto the cells, and the cells were fixed or imaged 30–36 hours post-transfection.

For the siRNA experiment, 6 or 9 μl of 20 μM siRNA and 4 μl of lipofectamine RNAi MAX (Invitrogen, 13778150) were suspended in 100 μl of water (Sigma, W4502) and were incubated for 5 min in parallel, then mixed and incubated for another 15 min. This mixture was added to the 2.5 ml medium containing around 100,000 cells per well. Cells were then grown for specified hours before fixation and immunostaining or live-imaging analysis.

For the generation of HeLa Kyoto cell lines stably expressing AcGFP-Ect2^r^ or AcGFP-Ect2^r^_Δmem_, cells were cultured in a 10-cm dish at 80% confluency. These cells were then transfected with 6 μg of pIRES-AcGFP-FLAG-Ect2^r^ or pIRES-AcGFP-FLAG-Ect2^r^_Δmem_ plasmid using 12 μl of lipofectamine 2000 (Life Technologies, 11668019). After 36 hr, 400 ng/μl puromycin media was added for the selection. Isolated colonies were cultured, and clones were confirmed by immunostaining and immunoblot analysis.

### Plasmids and siRNAs

All human Ect2 clones were generated using cDNA from the human cells as a template via PCR amplification. siRNA resistant Ect2 full-length and Ect2^r^_Δmem_ was cloned into a pIRES-AcGFP-FLAG plasmid (a gift from Mark Petronczki) using Age1 and EcoR1 sites. NuMA full-length was cloned into the pIRES-AcGFP-FLAG plasmid using Age1 and EcoR1 sites as reported in Rajeevan et al., 2020. All the clones were confirmed by sequencing analysis.

### Drug treatments

HeLa cells were synchronized in anaphase by double thymidine release. Briefly, the cells were treated with 2 mM thymidine (Sigma-Aldrich, T1895) for 17 hr, released for 8 hr, followed by another round of thymidine treatment for 17 hr. 10.5 hr after 2nd thymidine release, the drug of choice was added onto the cells, as mentioned in the respective figure panels. To inactivate Aurora B kinase, cells were treated with ZM447439 (Selleckchem, S1103). For initiating premature anaphase entry, cells were treated with 10 µM RO-3306 (Selleckchem, S7747) for 5 minutes. To inhibit myosin II, cells were treated with 100 µM para-nitroblebbistatin (PNBB; Optopharma Ltd., DR-A-081) for 30 min or 12 hr. Following drug treatments, the cells were fixed and immunostained.

For immunoprecipitation analysis, HeLa Kyoto cells stably expressing AcGFP-Ect2^r^ were synchronized in prometaphase with 100 nM nocodazole for 17 hr, and released in 10 µM MG132 (Selleckchem, S2619) for 4 hr to obtain metaphase synchronized cell population. After that, cells were treated with 10 µM of Cdk1 inhibitor RO-3306 (Selleckchem, S7747) for synchronizing them in the anaphase-like state, as reported previously in Keshri et al., 2020.

### Indirect immunofluorescence and live-imaging

For immunofluorescence analysis, cells were fixed with cold methanol for 10 min and washed in PBST (PBS containing 0.05% Triton X-100) or fixed with ice-cold 10% Trichloroacetic acid (TCA, Sigma, T0699) for 15 minutes at 4°C and washed in PBST. Cells were blocked in 1% bovine serum albumin (Himedia; MB083) for 1 hr, followed by incubation with primary antibody for 4 hr at room temperature. The primary antibodies used were 1:200 rabbit anti-NuMA (Santa Cruz, sc-48773), 1:300 mouse anti-NuMA (Santa Cruz, sc-365532), 1:200 mouse anti-p150^Glued^ (Transduction Laboratories, 612709), 1:2000 mouse anti-α-tubulin (Thermo Scientific), 1:20,000 rabbit anti-GFP (Invitrogen, A11122), 1:1000 mouse anti-GFP (DSHB), 1:200 rabbit anti-Ect2 (Merck, 07-1364), 1:200 rabbit anti-Cyk4 (Bethyl, A302-797A), 1:200 rabbit anti-Mklp1 (Santacruz, sc-867), 1:200 mouse anti-RhoA (Santacruz, sc-418), 1:200 rabbit anti-Prc1 (Santacruz, sc-8356), anti-Aurora B (BD bioscience, AB_398396). After incubating with primary antibody, washes were done three times, 5 min each with PBST, following which cells were incubated with secondary antibody for an hour. After three washes with PBST, cells were incubated with 1 μg/ml Hoechst 33342 (Sigma-Aldrich, B2261) for 5 min. Following three washes with PBST, the coverslips were mounted using Fluoromount (SouthernBiotech; 0100-01). Secondary antibodies used were 1:500 Alexa Fluor 488 goat anti-mouse (Invitrogen, A11001), 1:500 Alexa Fluor 488 goat anti-rabbit (Invitrogen, A11008), 1:500 Alexa Fluor 568 goat anti-mouse (Invitrogen, A11004), and 1:500 Alexa Fluor 568 goat anti-rabbit (Invitrogen, A11011). Confocal images were acquired on an Olympus FV 3000 confocal laser scanning microscope using a 60X (NA 1.4) oil immersion objective. All the images were processed using ImageJ.

Time-lapse recording was performed on an Olympus FV 3000 confocal laser scanning microscope using a 40X (NA 1.3) oil immersion objective (Olympus Corporation, Japan) using an imaging dish (Eppendorf, 0030740017) at 5% CO2, 37°C, 90% humidity maintained by a Tokai Hit STR Stage Top incubator. Images were acquired either at the interval of 1 min, 2 min, or 3 min with 9–11 optical sections (3 μm apart).

### Immunoblotting and immunoprecipitation

For western blotting analysis, HeLa cells synchronized with 100 nM nocadazole for 16– 20 hr were lysed in lysis buffer (50 mM Tris, pH 7.4, 150 mM NaCl, 2 mM EDTA, 2 mM EGTA, 25 mM sodium fluoride, 0.1 mM sodium orthovanadate, 0.1 mM phenylmethylsulfonyl fluoride, 0.2% Triton X-100, 0.3% NP-40, 100 nM Okadaic acid, and complete EDTA-free protease inhibitor) for 2 hours on ice and after a spin of 14,000 rpm, cell supernatant was denatured at 99°C in 2X SDS-PAGE buffer and analyzed by SDS-PAGE. After transfer to nitrocellulose membrane (Biorad, 1620115), blocking was done for an hour using 5 % skimmed milk (Himedia, GRM1254) in PBST. The membrane was then incubated with primary antibodies overnight at 4°C. The primary antibodies used were 1:1000 rabbit anti-Ect2 (Merck, 07-1364), 1:1000 rabbit anti-Cyk4 (Bethyl, A302-797A), 1:1000 rabbit anti-Mklp1 (Santacruz, sc-867), 1:1000 mouse Anillin (Santacruz, sc-271814), 1:5000 mouse anti-β-actin (Santa Cruz, sc-58673), 1:1000 rabbit anti-Prc1 (Santacruz, sc-8356), 1:1000 mouse anti-p150Glued (BD Bioscience, 612709); 1:20,000 rabbit anti-GFP(Invitrogen, A11122), 1:1000 rabbit anti-NuMA (Santa Cruz, sc-48773). After three washes in PBST (0.05% Tween-20; P1379; Sigma) for 5 min each, the membrane was incubated with HRP-conjugated goat anti-rabbit IgG (Jackson ImmunoResearch Laboratories INC., 111-035-045) and HRP-conjugated goat anti-mouse IgG (Bethyl, A90-116P) for an hour at room temperature. The membrane was further washed three times for 5 min each in PBST, and developed using Luminsol (Merck, WBLUF0500).

For immunoprecipitation, the harvested cells were lysed in lysis buffer [10 mM Tris, Ph7.5, 150 mM NaCl, 0.5 mM EDTA, 1 mM PMSF(Calbiochem, 7110), 0.5% NP-40 and Complete EDTA-free protease inhibitor (Merck, 539134)]. 2mg equivalent of cell lysate was incubated with 30 µl of GFP-Trap agarose beads (Chromotek, ACT-CM-GFA0050) at 4°C for 2 hours. After binding, the beads were washed two times with wash buffer (Lysis buffer without 0.5% NP-40) at 4°C. The bead-bound complex was denatured at 99°C in 2X SDS buffer and was analyzed by immunoblotting.

### Quantifications and statistical analysis

All quantifications were done in ImageJ. Quantification of membrane signal of NuMA or GFP intensity (Figure 1Q and 1T; Figure 2D; Figure 4J) was measured by calculating the ratio of the mean intensity of equatorial membrane signal (of a rectangular region of interest of area 3.5 µm^2^) divided by the mean intensity value in the cytoplasm (similar area) and correcting for the background signal (an analogous area outside the cell).

Quantification of membrane AcGFP-Ect2^r^ (Figure 3L) and membrane RhoA (Figure 5J) intensity was measured by calculating the ratio of the mean intensity of equatorial membrane signal (of a rectangular region of interest of area 3.5 µm2) divided by the mean intensity of polar membrane signal (similar area) and corrected for background signal.

The linescan intensity profile was measured using Image J. Briefly, the ‘straight line’ tool was used to draw a line of 2.5 µm through the cortex of the anaphase cells to obtain the line scan plot intensity values for NuMA at the equatorial membrane (Figure 1L-1P), and equatorial and polar membrane for RhoA (Figure 5G and 5H)

The cell length (Figure 2C-D) and cytokinetic furrow width (Figure 6F-6J) were measured at the mid-plane of the DIC live imaging movies using the line tool in ImageJ. The freehand tool on ImageJ was used for accurately outlining the cell cortex to get the linescan plot of cortical NuMA and RhoA (Figure 1A-1C), and AcGFP-NuMA and AcGFP-Ect2^r^ at t=12 minutes after anaphase onset (Figure 3M-3O). The freehand tool of ImageJ was used to determine the RhoA zone in anaphase cells by outlining the RhoA enriched equatorial membrane, and the whole cell perimeter (Figure S5D-S5E). RhoA zone (%) was determined by the equation mentioned in Figure S5D.

The inter-chromatid distance (Figure S4B) was quantified in maximum intensity projected live imaging movies of cells expressing AcGFP-NuMA;mCherry-H2B every 1 min after anaphase onset using ImageJ.

The relative astral microtubule intensity of anaphase cell was measured using imageJ. Briefly, maximum intensity projected images were used to calculate the total (astral and spindle) microtubule intensity (I^n^_total_) and the spindle microtubule intensity (I^n^_spindle_) of the cell by using the selected area shown in Figure S5F. The relative astral microtubule intensity was further calculated using the equation shown in Figure S5F.

To calculate the significance of the differences between two mean values, two-tail Student’s t-tests were performed. The p-value was considered to be significant if p < 1.5 using GraphPad Prism 8. The significance are mentioned as ns-p≥0.05; *-p<0.05; **-p<0.01; ***-p<0.001).

## Acknowledgements

We thank Daniel Gerlich, Mark Petronczki, and Anthony Hyman (MPI-CBG, Dresden) for providing plasmids and cell lines. We thank Arshad Desai, Karen Oegema, Andrew Goryachev, and Alissa Schlientz for fruitful discussion during this study, and their critical comments on the manuscript. We further thank Iain Hagan, Emanuelle Derivery, and the members of SK lab for their valuable suggestions on the manuscript. We thank Sukriti Kapoor for help with the working model. We thank DST-FIST, UGC Centre for the Advanced Study, DBT-IISc Partnership Program and IISc for the infrastructure support. This work is supported by the Department of Biotechnology (DBT)-Indian Institute of Science Partnership Program, DBT grant (<u>BT/PR36084/BRB/10/1857/2020</u>), and by grants from the Wellcome Trust/DBT India Alliance Fellowship (IA/I/15/2/502077 to S. Kotak). S. Kotak is a Wellcome Trust DBT-India Alliance Intermediate Fellow.

## AUTHOR CONTRIBUTIONS

Conceptualization: S.K.; Methodology: S.S. and A.R.; Validation: S.S., A.R., and S.K.; Formal analysis: S.S., A.R., and S.K.; Investigation: S.S., A.R., and S.K; Resources: S.K.; Data curation: S.S., A.R., and S.K.; Writing - original draft: S.K.; Writing - review & editing: S.S., A.R., and S.K.; Supervision: S.K.; Project administration: S.K.; Funding acquisition: S.K.

---

**Supplemental Figure 1.**
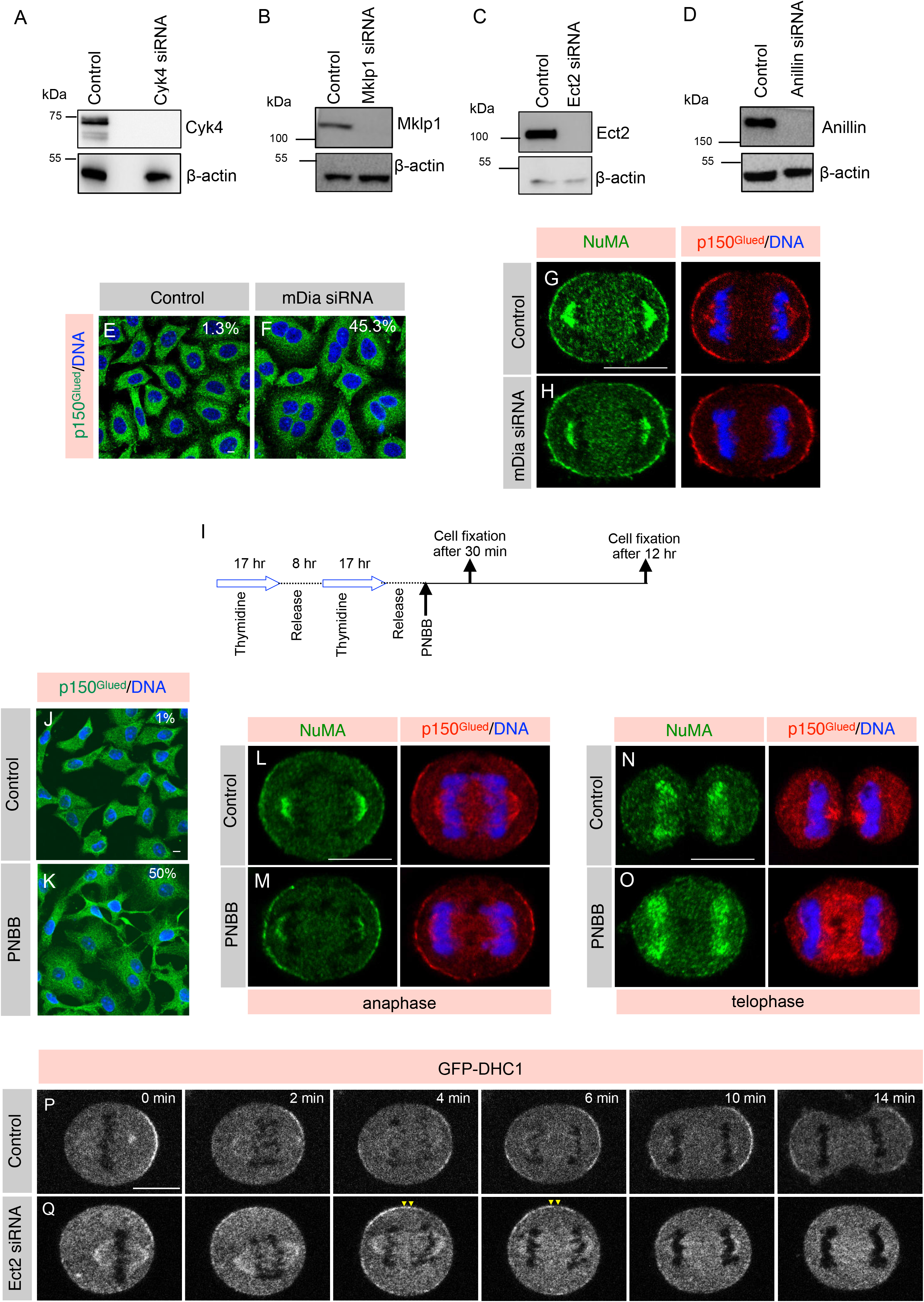
Formin and Myosin II activity is not critical for NuMA exclusion. (A-D) Assessing the depletion efficiency of siRNA against Cyk4, Mklp1, Ect2, or Anillin by immunoblot analysis. Mitotically synchronized protein extracts made from HeLa Kyoto cells that were either transfected with control siRNA, or siRNA-against above mentioned genes for 60 hr. Resulting blot was probed with antibodies directed against Cyk4 (A), Mklp1 (B), Ect2 (C), or Anillin (D). Antibodies against β-actin were used for loading control. In this and other Figure panels for immunoblot analysis, the molecular mass is indicated in kilodaltons (kDa) and is shown on the left. (E, F) Immunofluorescence (IF) analysis of HeLa cells that are either transfected with control siRNA (E) or siRNA-against mDia (Formin; F). Cells were fixed after 60 hr of siRNA transfection and stained with anti-p150^Glued^ antibodies (green). In this, and other IF-analysis panels DNA is shown in blue. Multinucleation percentage (%) is shown in the Figure panel. (n>500 cells were analyzed from two independent experiments). Scale bar in this panel and following panels represent 10 µm. (G, H) HeLa cells transfected with control siRNA (G) or siRNA against mDia (H). Cells were fixed after 60 hr of siRNA transfection and thereafter costained with anti-NuMA (green), and anti-p150^Glued^ (red) antibodies. (I) Synchronization scheme for myosin II inactivation using para-nitroblebbistatin (PNBB) to analyze cells in anaphase for NuMA and p150^Glued^ localization, and after 12 hr for assessing cytokinesis failure. (J, K) IF analysis of mitotically synchronized populations of HeLa cells as mentioned above were either treated with DMSO (Control; J) or PNBB (K) for 12 hr. Thereafter, cells were fixed and stained with anti-p150^Glued^ antibodies (green). Multinucleation percentage (%) is shown in the Figure panel. (n>500 cells were analyzed, from two independent experiments). (L-O) HeLa cells during early anaphase (L, M) or in telophase (N, O) that were either treated with DMSO (Control) or PNBB for 30 min. Cells were fixed and costained with anti-NuMA (green), and anti-p150^Glued^ (red) antibodies. Please note that during telophase (N, O), NuMA has localized to the nucleus, but the cleavage furrow has not formed because of myosin II inactivation. n>50 cells, from two independent experiments, and the representative cell is shown here. (P, Q) Confocal live-imaging analysis of HeLa Kyoto cells stably expressing GFP-DHC1 that were transfected with control siRNA (P) or siRNA against Ect2 (Q). Recording was started 30 hr post transfection for control and Ect2 siRNA. ‘0 min’ time-point represent metaphase to anaphase transition. Yellow arrowheads depict GFP-DHC1 localization at the equatorial membrane. Representative images from the time-lapse recording are shown here. (n>10 cells).

**Supplemental Figure 2.**
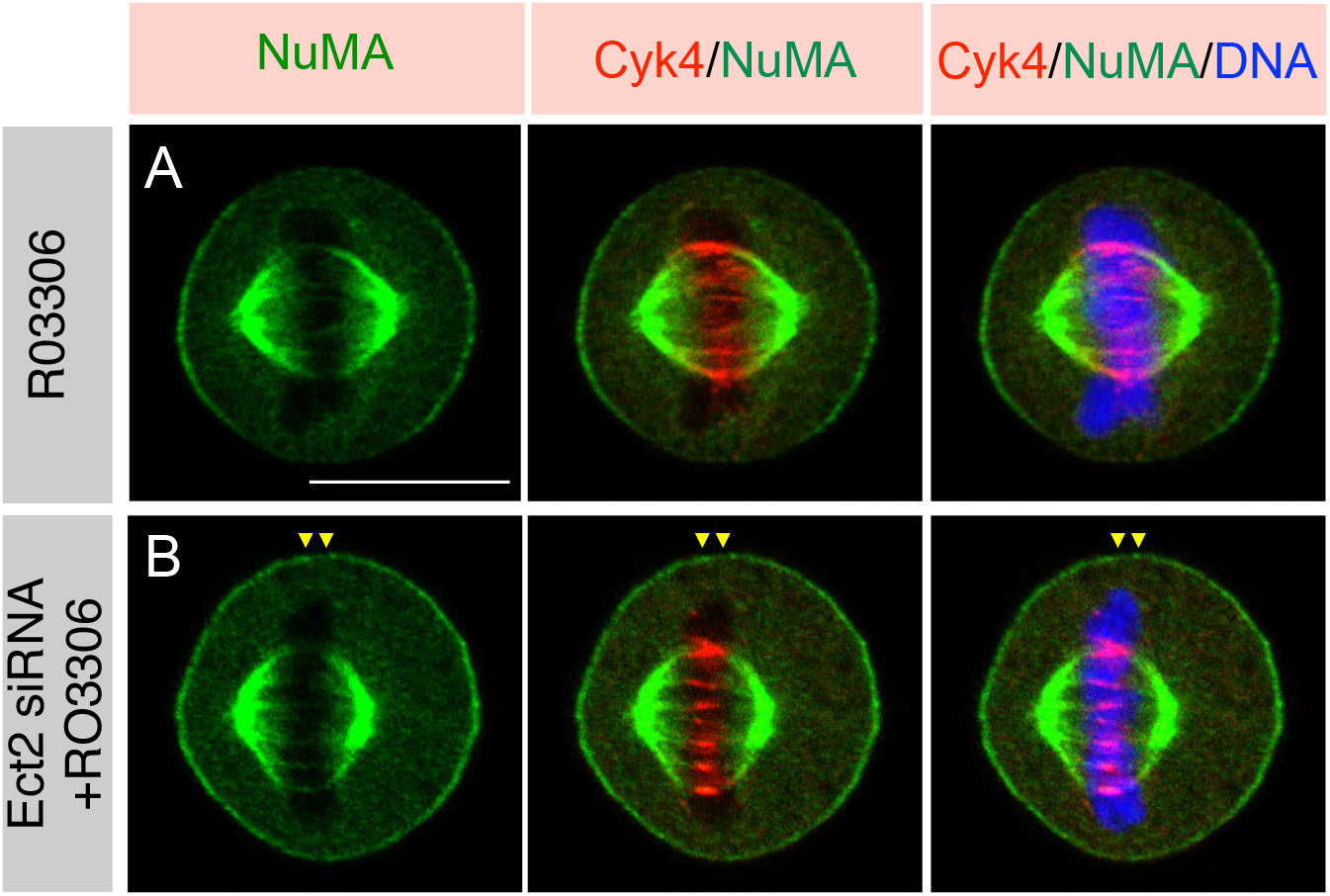
NuMA exclusion from the equatorial membrane is not dependent on cell length. (A, B) IF analysis of HeLa cells that were either control siRNA transfected and treated with RO-3306 (A), or transfected with siRNA-against Ect2 and treated with RO-3306 (B) for 5 min. Cells were fixed and costained using anti-NuMA (green), and anti-Cyk4 (red) antibodies. Yellow arrowheads depict NuMA localization at the equatorial membrane. Scale bar represent 10 µm. More than 30 cells were analyzed and the representative cells are shown here.

**Supplemental Figure 3.**
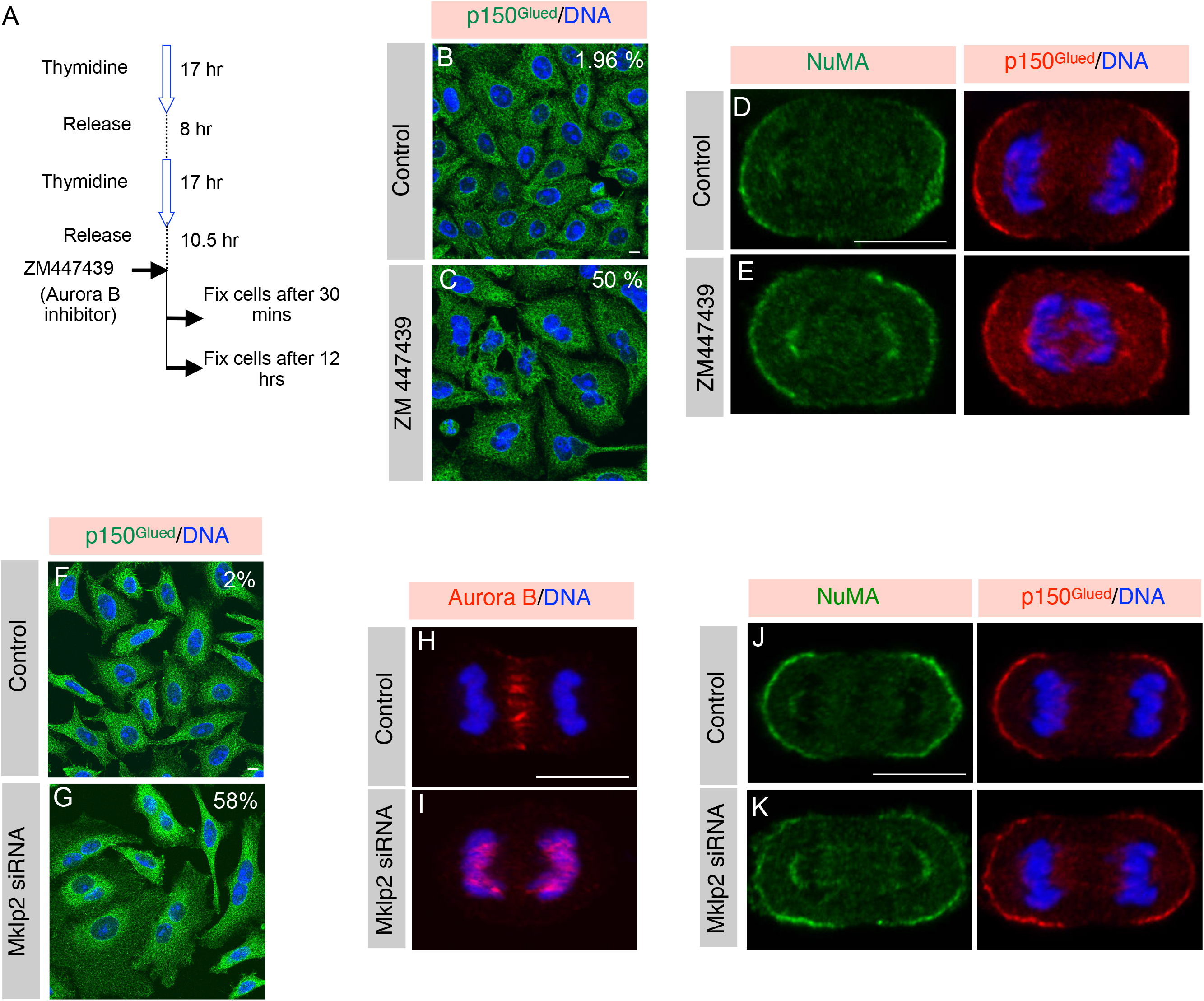
Central spindle localized Aurora B is not essential for NuMA exclusion from the equatorial membrane. (A) Synchronization method for inactivation of Aurora B using ZM447439 to analyze cells in anaphase for NuMA and p150^Glued^ localization, and after 12 hr for assessing cytokinesis failure. (B, C) IF analysis of mitotically synchronized populations of HeLa cells as mentioned above either treated with DMSO (Control; B) or ZM447439 (C) for 12 hr. Thereafter, cells were fixed and stained with anti-p150^Glued^ antibodies (green). Scale bar in the panel and following panels represent 10 µm. Multinucleation percentage (%) is shown in the figure panel. (n>500 cells were analyzed each from two independent experiments). (D, E) Mitotically synchronized populations of HeLa cells were either treated with DMSO (Control; D) or ZM447439 (E) for 30 min. Cells were fixed and costained with anti-NuMA (green), and anti-p150^Glued^ (red) antibodies. More than 50 anaphase cells were analyzed from two independent experiments, and the representative cell is shown here. (F, G) IF analysis of HeLa cells in interphase that were either transfected with control siRNA (F) or siRNA-against Mklp2 (G). Cells were fixed after 40 hr of siRNA transfection and stained with anti-p150^Glued^ antibodies (green). Multinucleation percentage (%) is shown for each siRNA condition. (n>500 cells from two independent experiments). (H, I) IF analysis of HeLa cells during anaphase that were either transfected with control siRNA (H) or siRNA-against Mklp2 (I). Cells were fixed after 40 hr of siRNAs transfection and thereafter stained with anti-Aurora B antibody (red). (J, K) IF analysis of HeLa cells in anaphase that were either transfected with control siRNA (Control; J) or siRNA-against Mklp2 (K). Cells were fixed after 40 hr of siRNA transfection and costained with anti-NuMA (green) and anti-p150^Glued^ (red) antibodies. More than 50 anaphase cells were analyzed from two independent experiments, and the representative cells are shown here.

**Supplemental Figure 4.**
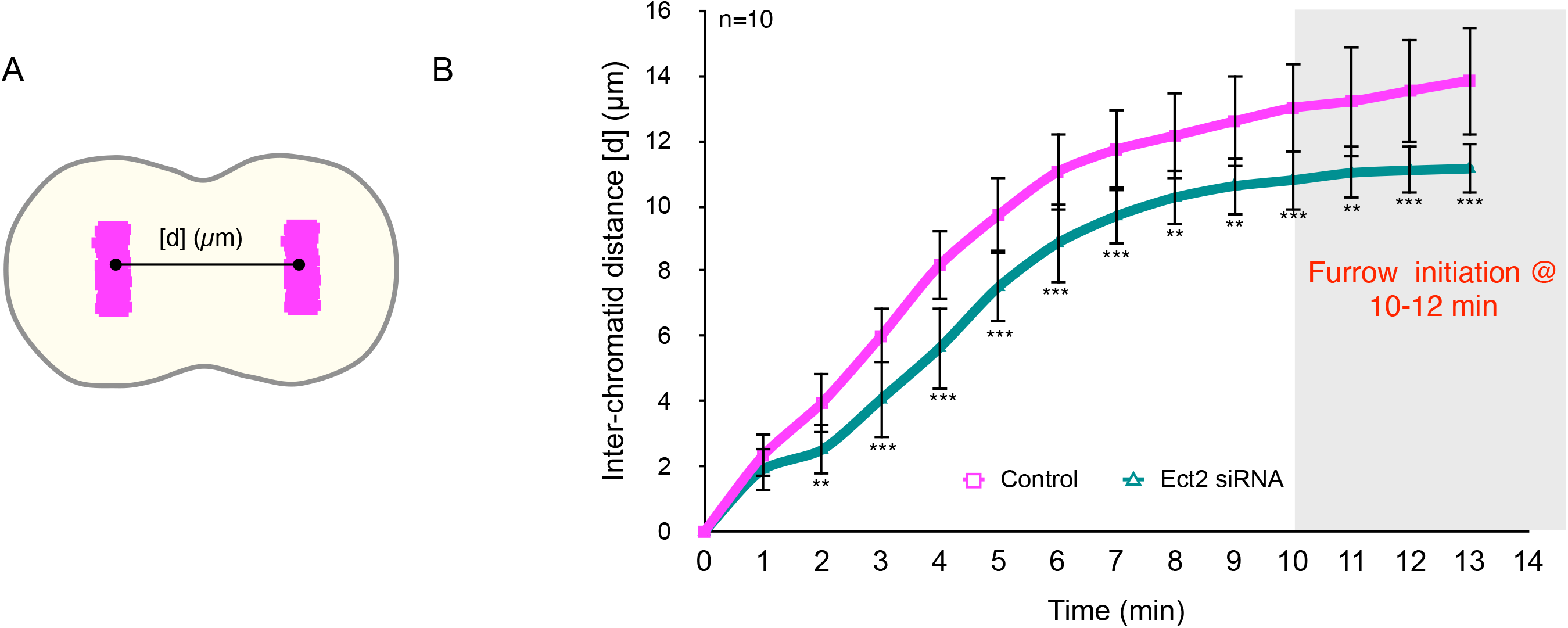
Equatorial localization of NuMA-based complexes impact proper chromosomes separation in anaphase. (A, B) Schematic representation for the calculation of the distance [d] between inter chromatids in cells undergoing anaphase progression at a various time interval (A). Quantification of inter chromatids distance in HeLa Kyoto cells that are stably coexpressing AcGFP-NuMA and mCherry-H2B and are transfected with either control siRNA or siRNA-against Ect2 (B) (n = 10 cells from two independent experiments as depicted in the figure panel; see Experimental procedures. error bars: SD).

**Supplemental Figure 5.**
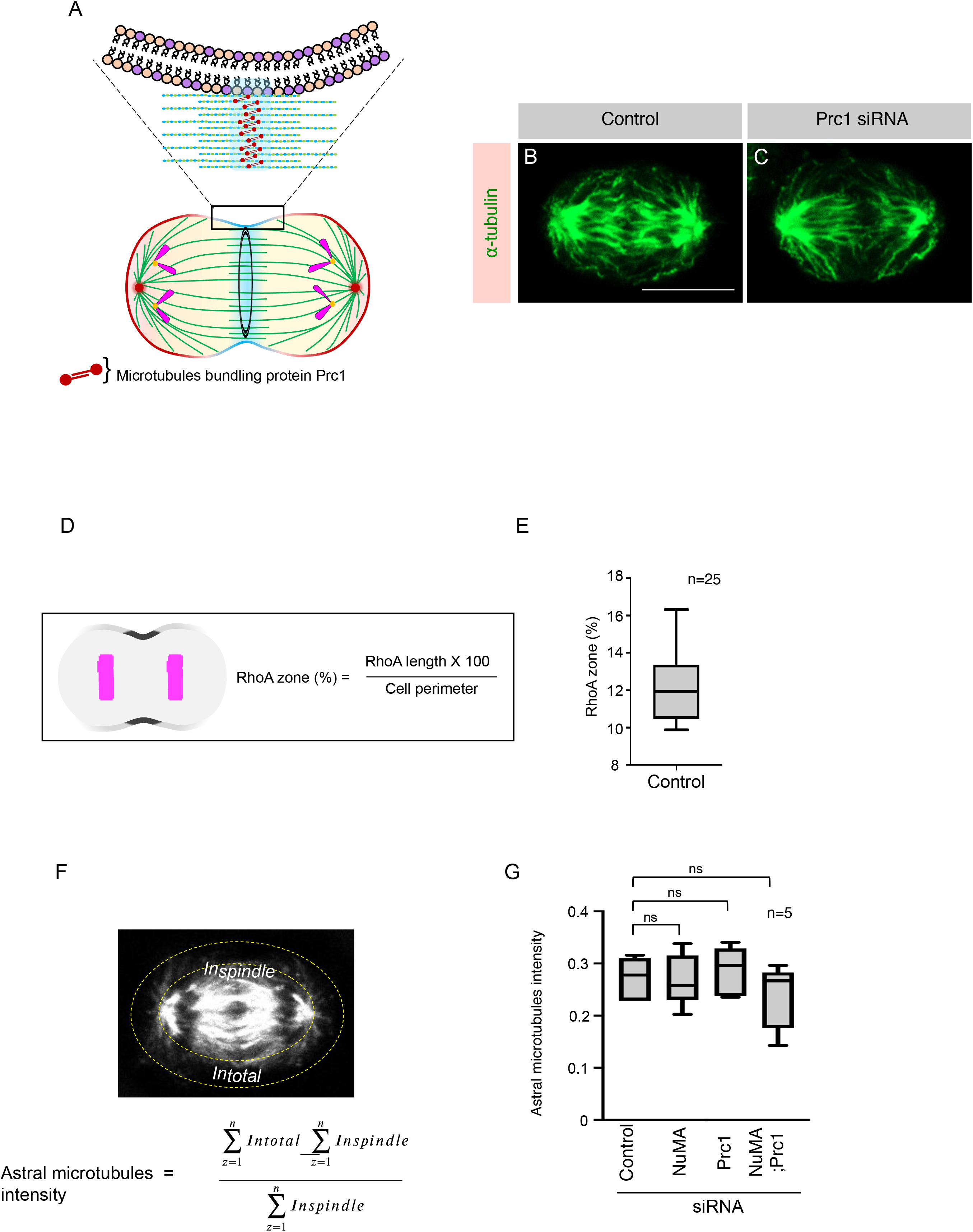
Astral microtubules are not affected in cells codepleted for Prc1 and NuMA. (A) A schematic representation of anaphase cells and highlighting the midzone microtubules bundles created by Prc1 (Protein Regulator of Cytokinesis 1) homodimer. (B, C) IF analysis of HeLa cells that were either transfected with control siRNA (B), or siRNA directed against Prc1 (C) for 36 hr were fixed, and stained with antibodies directed against ⍺-tubulin (in green). Scale bar in the panel and following panels represent 10 µm. (D, E) Schematic representation of quantification method (D) used to determine the RhoA zone (in grey), and outcome of such analysis in cells transfected with control siRNA (E). (n=25 cells; error bars: SD). (F, G) Representative IF image of HeLa Kyoto cell in anaphase fixed and stained with α-tubulin antibody (grey). The areas used to analyze relative astral microtubules intensity is shown (F). Quantification of relative astral microtubule intensity in cells transfected with control siRNA (control), Prc1 siRNA, NuMA siRNA or NuMA and Prc1 siRNA (G).The intensities of the spindle (I_spindle_) and the total cell (I_total_) were determined with ImageJ software. Relative astral MT intensity (Iastral, rel) was calculated by [(Itotal-Ispindle)/ Ispindle]. (n = 5 cells in each condition. p>0.1 in each condition w.r.t control; error bars: SD).

